# Climatic factors and host species composition at hibernation sites drive the incidence of bat fungal disease

**DOI:** 10.1101/2023.02.27.529820

**Authors:** AS Blomberg, TM Lilley, M Fritze, SJ Puechmaille

## Abstract

Emerging infectious diseases pose a serious threat to wildlife, and their occurrence will likely be further exacerbated due to climate change. The aim of our study was to investigate whether the occurrence of White-nose disease (WND), a fungal disease of hibernating bats, can be predicted using local climatic conditions and host species abundance at hibernation sites. In addition, we used our model to predict areas potentially at risk if the pathogen is introduced and investigated how the potential distribution of WND may shift in the future due to climate change. We employed logistic regression as part of our ecological niche modelling approach, integrating climate and census data as explanatory variables, along with WND status (the response variable) obtained from 448 hibernacula. This approach allowed us to predict regions at elevated risk of WND by applying these climatic variables to current global climatic conditions and a climate change scenario. A model incorporating data on mean annual surface temperature, precipitation, and three host species reliably predicted WND occurrence in Europe. This model demonstrated robust transferability beyond Europe, as confirmed by both theoretical and empirical assessments (e.g., accurately predicting the observed mortality events in North America). We identified several high-risk areas in the southern hemisphere and demonstrate that climate change may cause a remarkable shift in the distribution range of WND. Our results highlight the importance of environmental factors in controlling the manifestation of disease in localities where both the pathogen and suitable hosts are present. We pinpoint several areas requiring increased surveillance and precautions to avoid the introduction of the pathogen, and show that climate change has massive potential to reshape and expand these areas, putting new populations at risk.

## 1 INTRODUCTION

The emergence and re-emergence of infectious diseases has increased globally during the past decades (Daszak 2000). Given that climatic factors, such as rainfall and temperature, contribute significantly to disease dynamics and host abundance (Dabaro 2021, Aune et al. 2021, Xu et al. 2023) it is likely that climate change further exacerbates the incidence of these diseases. Fungal pathogens have emerged as a major threat to humans, wildlife, and plants (Fisher et al. 2012, 2016), with *Candida auris* often causing severe candidiasis in humans, and *Batrachochytrium dendrobatidis* being responsible for the population crashes of several amphibian species globally (Bosch et al. 2007, Garcia-Solache and Casadevall 2010, Casadevall et al. 2019, Nnadi and Carter 2021). Therefore, developing new, effective, and minimally invasive monitoring systems and prediction tools are crucial for disease surveillance and conservation. Information regarding the distribution of pathogens or diseases, both current and future, is of paramount importance, yet it is often lacking on a global scale. Bridging these knowledge gaps can be achieved through the use of modelling techniques that can provide such information.

Ecological niche modelling is widely used in ecology to predict the potential geographic distribution of species in light of their environmental limitations. It is rapidly emerging as the preferred method for disease risk mapping, establishing itself as the gold standard in the field (Johnson et al. 2019). The most popular approaches used can be positioned on a continuum, delineated by the level of explicit representation of underlying processes (Dormann et al. 2012). On one end of the continuum, correlative models infer parameters by examining the correlations between current species distributions and environmental factors while on the other end of the continuum, mechanistic models explicitly integrate causal relationships between environmental conditions and the performance of an organism. Correlative models require the user to gather sufficient presence data for the species under investigation, along with a set of environmental variables influencing the species’ distribution. In contrast, mechanistic models necessitate that the user identifies critical constraints on species survival and fecundity, such as physiological requirements, and develops a parameterized mechanistic model to link these constraints. Identifying key constraints and parametrizing the model typically demands extensive empirical studies, which are available for only a very limited number of species (Elith 2017). Furthermore, parameterization is particularly critical as errors in parameters can compound leading to poor accuracy in prediction (Buckley et al. 2010). These challenges, associated with models on the mechanistic end of the continuum, contribute to less frequent use compared to correlative models. Despite the different foundations and limitations of approaches at the continuum’s extremes (Dormann et al. 2012), they often provide congruent outcomes when compared, including under climate change scenarios (e.g., Buckley et al. 2010, Kearney et al. 2010, Elith 2017).

We herein aim to use ecological niche modelling to better understand the regions of the world that have environmental conditions conducive for the White-nose disease (WND). WND, associated with the white-nose syndrome, is an emerging fungal disease in bats (Whiting-Fawcett et al. 2025). It is caused by *Pseudogymnoascus destructans*, a psychrophilic fungus infecting bats during hibernation (Blehert et al. 2009, Gargas et al. 2009, Frick et al. 2016). The fungus is native to Eurasia where it has coexisted with bats for a considerable time (Leopardi et al. 2015, Drees et al. 2017, Fischer et al. 2025) and bats adapted to *P. destructans* by developing immunological tolerance (Harazim et al. 2018, Lilley et al. 2019, Fritze et al. 2021a, Whiting-Fawcett et al. 2021, 2025). Following the recent introduction of its causative agent, WND has caused severe depletion to populations of several North American bat species (Thogmartin et al. 2012, Frick et al. 2015). This resulted in a highly variable outcome, ranging from no or limited apparent negative effects on the host, as observed in the native range in Eurasia, to host mortality of several species in the introduced North American range (reviewed in Whiting-Fawcett et al. 2025; Fig 1). Despite yielding divergent outcomes in this host-pathogen interaction, WND, diagnosed through skin erosion induced by *P. destructans*, has been identified across Eurasia and North America (Zukal et al. 2016, Martínková et al. 2018).

**Figure 1.**
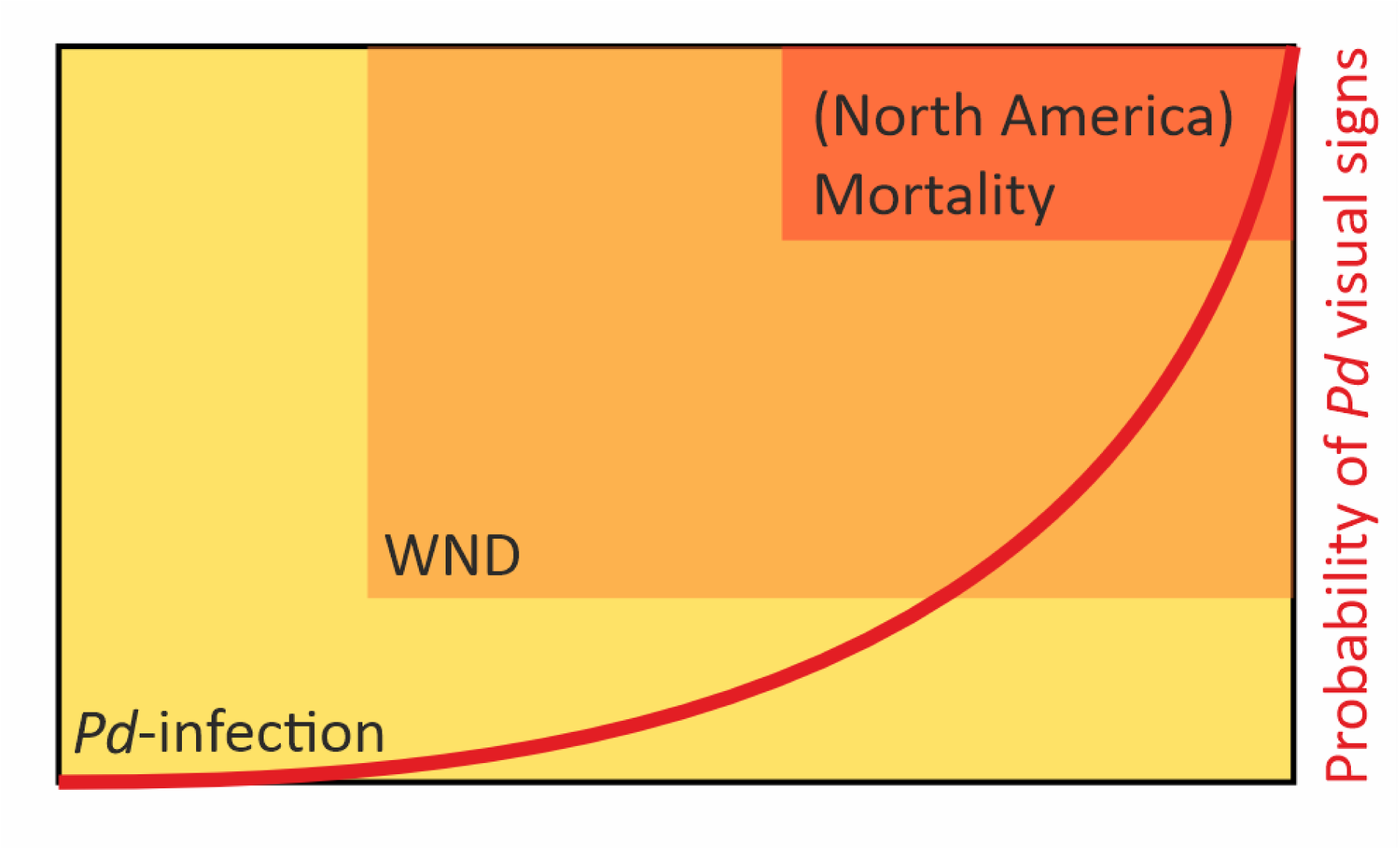
Schematic representation of the link between *Pd*-infection (*Pd*-load increasing from left to right), White-Nose Disease (severity increasing from left to right) and the resulting mortality in relation to the probability of visual detection of *P. destructans* (*Pd*) on live bat hosts (box sizes not proportional to incidence) as predicted from existing literature (Janicki et al. 2015, Hoyt et al. 2021, Fritze et al. 2021a, 2021b).

Similar to other diseases, WND is manifested when a pathogen and a susceptible host co-occur in a suitable environment (Scholthof 2007). Therefore, as in most other systems, disease is not manifested throughout the distribution range of the pathogen, if susceptible hosts are not present, or when environmental conditions are not suitable. Despite important efforts to better understand the disease over the past 15 years, building a global and reliable mechanistic model for such a complex multi-host system is currently not possible and is also unlikely to be in the foreseeable future (Puechmaille et al. 2011a, Hoyt et al. 2021, Whiting-Fawcett et al. 2025). That being said, several key processes have been identified and provide critical knowledge to inform variable selection for models towards the correlation spectrum. We aim to use this knowledge to include only those abiotic variables in our modelling framework for which we have strong evidence of causal involvement, as these relationships are likely to remain stable across spatiotemporal scales. One of these key factors is temperature; it has been demonstrated under experimental laboratory conditions that *P. destructans* optimally grows on cultures at around 12.5–15.8 °C, with no detectable growth below zero or above 20 °C (Chaturvedi et al. 2010, Verant et al. 2012). Hence, temperature is a key variable shaping the potential distribution of the pathogen, and hence the disease. Similarly to temperature, *P. destructans* growth and spore production are maximal at high air humidity (Marroquin et al. 2017). Given its psychrophilic nature, and its slow growth, *P. destructans* is thought to be restricted to growth on underground hibernating bats, which typically hibernate at temperatures of 0–15 °C and in conditions of high air humidity (Webb et al. 1996). These conditions favour extended torpor of bats, as low ambient temperatures promote longer bouts of torpor, resulting in reduced energetic costs and lower evaporative water loss. This effect is further enhanced by the high humidity in hibernacula (Crowley et al. 2026). Extended periods of torpor provide opportunities for fungal growth on the bat host. Temperatures inside underground hibernation sites correlate with outside mean annual surface temperatures (MAST) (Dwyer 1971, Martínková et al. 2018, McClure et al. 2020), and a similar correlation is evidenced between underground humidity levels and annual precipitations (Perry 2013). Therefore, as a result of the mechanistic relationship between temperature, torpor patterns, and the growth of *P. destructans* (and, consequently, infection intensity), MAST and precipitation are expected to be major drivers and hence predictors of WND. The ongoing and future changes in MAST expected through climate change will therefore modify the areas climatically suitable for WND occurrence.

Regarding susceptible hosts, closely related species tend to share ecological, physiological, and immunological traits, and are generally also more likely to share parasites (Huang et al. 2014, Clark and Clegg 2017, Clark et al. 2018). In the case of *P. destructans* infection, closely related host species are most likely to be infected (Zukal et al. 2014), with species in the genus *Myotis* being most affected. That being said, based on the list of 61 species currently known to be affected by WND or *P. destructans* infection across Eurasia and North America (Zukal et al. 2016, Hoyt et al. 2021), it is clear that the fungus is neither species-, genus- nor family-specific. In fact, species belonging to three families in each of the two suborders of bats, Yinpterochiroptera and Yangochiroptera, have been documented with *P. destructans* infection. A comparative study investigating host traits and their phylogeny revealed that neither ecological nor evolutionary constraints on hibernating bats pose a barrier to this generalist fungal pathogen (Zukal et al. 2014). Therefore, hibernation in contaminated hibernacula under favourable abiotic conditions appears to be the main factor influencing risk of infection (Zukal et al. 2014, Wu et al. 2025).

Although *P. destructans* is a generalist species, all hibernating bat species are not equally conducive of the disease (Hoyt et al. 2021, Whiting-Fawcett et al. 2025). The species most noticeably affected by WND in Europe is *Myotis myotis* (greater mouse-eared bat) (Puechmaille et al. 2011b), which has been recorded with the highest pathogen loads (Zukal et al. 2014), as well as the highest number and prevalence of lesions caused by the fungus, compared to other species sharing the same hibernacula (Zukal et al. 2014, 2016). On the other hand, *Rhinolophus* species typically harbour lower pathogen load compared to *Myotis* species while sharing similar hibernation conditions (Zukal et al. 2016, Hoyt et al. 2021). Therefore, at a more local scale, the presence or detection of the disease might also be affected by the host’s species present and their respective abundance (see also Laggan et al. 2023). If not taken into consideration, the identity and abundance of hosts could blur the relationship between abiotic factors and WND status. Thus, on a large scale, above surface climatic conditions significantly contribute to shaping the current geographical distribution of WND in Eurasia while species composition of the hosts seems to play a role at a more local scale. Climate change is expected to alter the MAST and hence microclimate conditions of hibernacula, placing new species and populations at risk, as the fungal pathogen extends its distribution to areas that have previously been outside its environmental growth conditions. In addition, the disease may begin to manifest in areas where the pathogen is currently present but prerequisites, such as suitable hosts or environmental conditions, are not met. Therefore, information on how climatic factors and host composition drive the occurrence of WND, and how climate change may contribute to the future spread of the disease, is vital for risk assessments and targeting monitoring as well as prevention/conservation measures.

Here, we leverage weather data and hibernation site census data from 448 European locations to model the effects of local climate (mean annual surface temperature, annual rainfall) and species abundance on WND occurrence. This ecological niche model then predicts regions that are climatically suitable for WND under current and future conditions. Our model identifies areas with suitable environmental conditions, but it does not predict the disease outcome (mortality, recovery, etc.) which is context dependent (Whiting-Fawcett et al. 2025).

## 2 MATERIAL AND METHODS

### 2.1 Data and data acquisition

The data consist of annual hibernation site censuses from 448 sites distributed across 34 European countries (Figure 2), obtained from those performing the censuses. Of the censused sites, parts of the data from 272 counts have previously been published by Fritze and Puechmaille (2018). The dataset includes species composition and species abundance in censused hibernacula, which was used here for the first time, and information on whether signs of WND (fungal growth on the wings and/or muzzles) have been visually observed on a live bat at the site during censuses (1 = yes, 0 = no; see also Fritze and Puechmaille 2018). Given that the visual observation of *P. destructans* on bats simultaneously provides information on fungal colonisation and wing damage, it is herein used as a proxy for the presence and severity of the disease (see Fritze et al. 2021b for more details; Horáček et al. 2014). This is distinct from the probability of encountering the fungus causing the disease, as measured by molecular methods (e.g. qPCR, LAMP; Muller et al. 2013, Niessen et al. 2022), which does not necessarily indicate the presence of the disease (Janicki et al. 2015, McGuire et al. 2016, Urbina et al. 2020, Fritze et al. 2021b). Fischer et al. (2025) identified two cryptic species within *P. destructans*, provisionally designated *Pd*-1 and *Pd*-2. However, because these taxa occur in overlapping geographic regions and have been reported to cause similar disease manifestations in subterranean environments characterised by comparable temperature and absolute humidity conditions, we did not distinguish between them in the present analyses and instead treated them as a single operational unit for both model input and output. Accordingly, our predictions should be interpreted as reflecting the combined distribution of *Pd*-1 and *Pd*-2. Nevertheless, if the two fungal species differ in their abiotic niche requirements, these predictions are likely to more closely represent *Pd*-1 than *Pd*-2, as Fischer et al. (2025) showed that *Pd*-1 accounts for approximately 95% of *P. destructans* infections in Europe. For censuses, in cases where several had been performed in the same site, we retained the survey with the highest number of bat species observed. If the number of species was equal for several censuses of the same site, we then retained the census with the highest number of individuals. All censuses were performed between the years 2009–2022, generally during regular surveys to avoid additional disturbance to the bats. Bats were counted mostly in situ, but in some cases, photographs were used to evaluate the number of individuals, especially in large clusters. At some sites, the surveyors also measured ambient temperature (N = 356 sites, one to several measurement/site) and relative humidity (N = 228 sites, one to several measurement/site; see Figure S4 for a map of locations). To facilitate the deployment of the survey to a maximum number of sites and hence improve geographical coverage, no specific equipment brand/model of thermometer or hydrometer was imposed to record temperature and humidity. The protocol for data collection is further detailed by Fritze and Puechmaille (2018). We also used the mean annual temperature and annual precipitation for each site. These climatic data were extracted from the worldclim version2.1 database of global interpolated climate data (Hijmans et al. 2005, Frick and Hijmans 2017) as detailed in Frick & Hijmans (2017).

**Figure 2.**
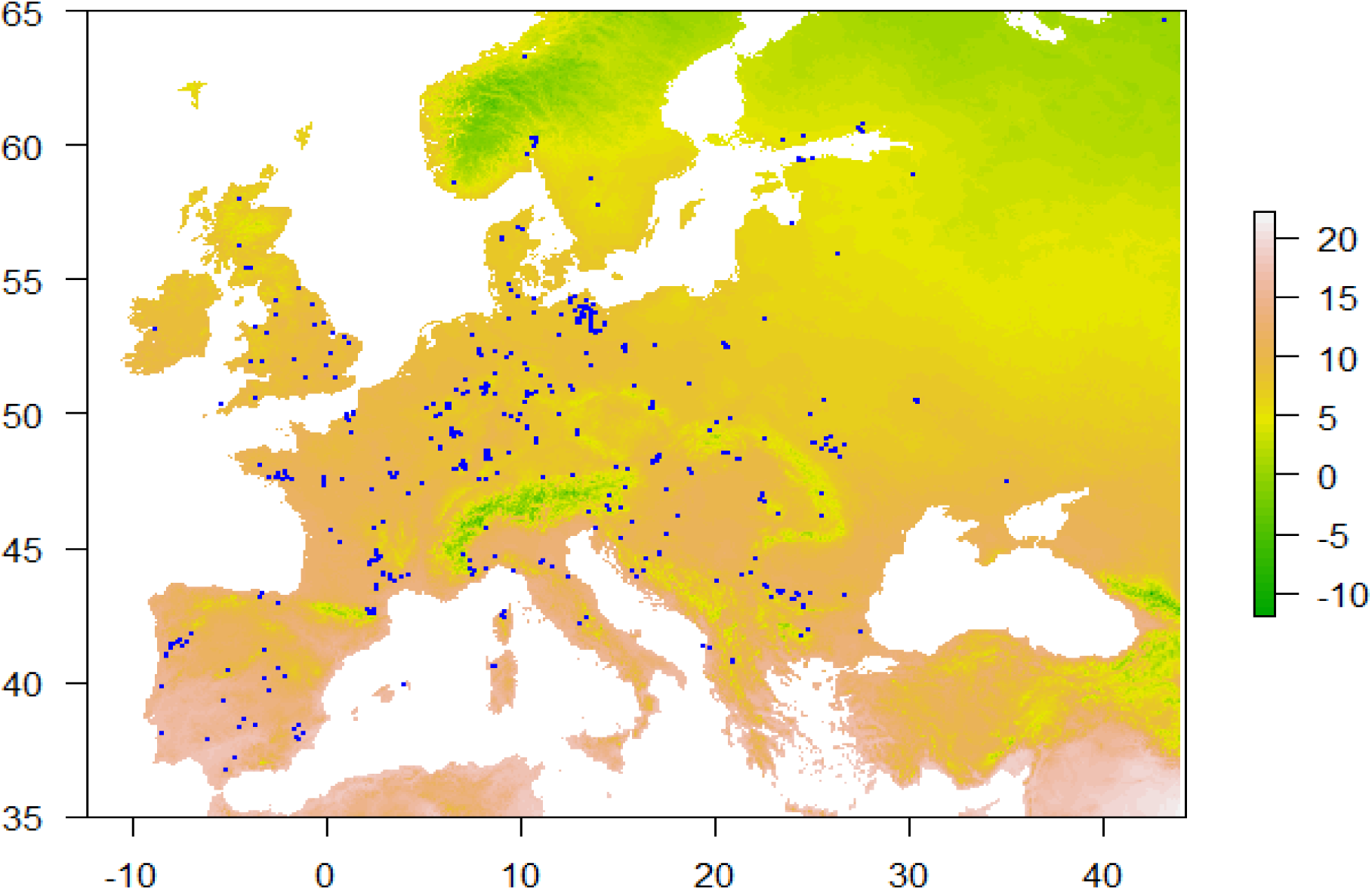
Map of observation sites with background colours representing Mean Annual Surface Temperature in °C.

In addition to the census and climatic data, we obtained both county-level WND associated mortality data, as well as data on the occurrence of the disease or the causative agent in North American hibernacula to be used for validation (see below). We extracted the mortality data from the USGS National Wildlife Health Center WHISPers event sharing database (Wildlife Health Sharing Partnership 2023), whereby we only selected events that were confirmed to be due to or associated with WND through histo-pathological evidence (i.e., cupping erosions *sensu* Meteyer et al. 2009). The data on WND occurrence and *P. destructans* presence was extracted from the White-nose syndrome spread map by the White Nose Syndrome Response Team (for data selection criteria please see statistical analysis below).

### 2.2 Statistical analysis

We fitted a logistic regression model (package stats in R (R Core Team 2023)) with a binomial distribution (“logit” link function) to investigate how climatic conditions (mean annual surface temperature (MAST) and annual rainfall at the site), species abundance, affect the probability of *P. destructans* having ever been visually observed on any living bat at the site (i.e., from now on, WND-suitability of the site). To decrease the number of considered explanatory variables, avoid overfitting the model, and diminish type I error (Burnham and Anderson 2002), we only included species that were present in at least 10 % of the censused hibernacula, as it is unlikely that rare species would play a key role in the WND-suitability at the scale herein investigated. We grouped species which are challenging to visually identify to species level during censuses (*Myotis myotis* and *M. blythii* (lesser mouse-eared bat) (262 sites, 60 148 individuals in total); *M. brandtii* (Brandt’s bat) and *M. mystacinus* (whiskered bat) (118 sites, 2 494 individuals in total); *M. crypticus* (cryptic myotis), *M. nattereri* (Natterer’s bat), and *M. escalerai* (Escalera’s bat) (179 sites, 15 168 individuals in total)). In addition to these three species groups, species included in the analysis were *M. daubentonii* (Daubenton’s bat) (221 sites, 35 604 individuals in total), *M. emarginatus* (Geoffroy’s bat) (63 sites, 2 948 individuals in total), *Barbastella barbastellus* (barbastelle bat) (53 sites, 4 737 individuals in total), *Plecotus auritus* (brown long-eared bat) (142 sites, 1 617 individuals in total), *Rhinolophus ferrumequinum* (Greater horseshoe bat) (153 sites, 19 078 individuals in total), and *R. hipposideros* (Lesser horseshoe bat) (145 sites, 7 694 individuals in total). We used the abundance of these species and species groups at each hibernation site, as well as MAST and precipitation, as explanatory variables in our analysis. We included MAST and annual precipitation in the model as quadratic terms, to account for the non-linear effect of those terms as determined from laboratory experiments (see introduction).

From the full logistic regression model with the thirteen above-mentioned explanatory variables, we selected the best model by using multi-model comparison (package MuMIn v 1.47.5 in R) and retained the model with lowest AICc. To ensure that our results were robust to the model selection approach, we also performed multi-model inferences via MuMIn. We obtained full model-averaged regression coefficients of the predictors from models within < 2, < 4 and < 7 AICc of the best model. So in total, we have five models: a full model, a best model, and three averaged models. We then used Wald Chi-square test to identify significant variables (considered p-values < 0.05) for each of the five models. We also calculated the confidence interval of regression coefficients, using the standard 95% CI corresponding to the 0.05 significance level and the 85% CI that is consistent with how terms are selected under the AIC-based model selection criteria (Arnold 2010, Sutherland et al. 2023). The results of these five models were nearly identical (Suppl. Figure S1 & Tables S1 to S5, S10). Hence, we opted to exclusively present the outcomes of the model with the lowest AICc (without employing model averaging) and used these findings for subsequent analyses, unless specified otherwise. All analyses were performed using R version 4.3.1 Patched (R Core Team 2023).

We checked for multicollinearity between the independent variables (species/species group abundances, MAST and precipitation) by using variance inflation factors (VIFs) calculated on the full model. For the species abundances, VIF values were < 3 for all species, suggesting that their regression coefficient estimates are reliable. As expected, MAST and precipitation strongly correlated with their respective quadratic terms (VIFs > 38 for all), potentially leading to model instability. We therefore investigated the robustness of our results by jackknifing each of our 448 data points (= sites) and recalculating the regression coefficient of the full model. The coefficient of variation of the jackknifed regression coefficients was lower than 1.79 % for the four variables with high VIFs, and the difference between the full model coefficients and the average coefficients of the jackknifed model was > 3,500 times smaller than their respective full model coefficients. These analyses indicate that the collinearity inherent to the inclusion of the quadratic terms does not create model instability. The results from the jackknifed analyses also confirm that the estimates for the species abundances are robust, except for *B. barbastellus* for which the standard deviation of the estimate is larger than its mean (See Suppl. Table S9). Nevertheless, given that *B. barbastellus* is unlikely to be an important species in relation to WND and is not significant in any models (see results), we did not further explore the variation between jackknifed datasets for that species.

We used Spearman’s correlation coefficient to test for correlation between MAST and temperature inside hibernacula (N = 356 sites, one measurement per site, Fig. S4). In addition, we investigated the corresponding hibernacula temperature to the optimal MAST for WND occurrence (derived from our logistic regression analysis described above), we also fit a linear model with the stats package in R (R Core Team 2023), with temperature inside hibernacula as the response variable and MAST as the explanatory variable. To investigate the relationship between annual precipitation and humidity inside hibernation sites, we first derived absolute humidity for each site from measures of relative humidity (N = 228 sites, one measurement per site; obtained during the censuses) by applying the formula presented by (Wagner and Pruß 2002). We then fit a generalised linear model with absolute humidity as the response variable, and precipitation as well as its quadratic term as explanatory variables to determine the absolute humidity corresponding to the optimal precipitation for WND occurrence.

### 2.3 Predictions

We used the coefficients derived from the best model of the logistic regression analysis to predict WND suitability for every cell (2.5-minutes resolution) of a raster map of the world. The model includes nine predictor variables, including MAST and precipitation, their respective quadratic term, and species compositions data (5 predictors; see Table 1). For MAST and precipitation, we used data obtained from the worldclim version2.1 database (Frick and Hijmans 2017), for recent climatic conditions (1970-2000) and future climatic conditions (2061-2080), the latter under model ACCESS-CM2, Shared Socio-economic Pathway 3-7.0. As no data exist across the world on species composition and species count at hibernacula, for every cell, we considered that every species was present and used the average species abundance calculated on the observed dataset. Therefore, except when species abundance is explicitly changed (see below), relative differences in the predictions are solely due to differences in climatic conditions. We performed all analyses using R version 4.3.1 Patched (R Core Team 2023). Additionally, to illustrate how the abundance of susceptible hosts affects the potential range of the disease, and to better capture the situation in North America when the disease emerged, we multiplied by ten the average number of *M. emarginatus*, the species with the strongest effect (see Results). The magnitude of the increase was established based on research conducted by (Frick et al. 2015), indicating that when the disease emerged, bat abundance at hibernacula in North America was tenfold higher than actual colony sizes in Europe. This 10-fold increase is therefore conservative as it is applied to a single species rather than across multiple species. Notably, even after multiplying the average number of *M. emarginatus* by ten, the resulting number is still approximately 11 times lower than the maximum observed count for that species (50 versus 550). Hence no data extrapolation is involved.

**Table 1.**
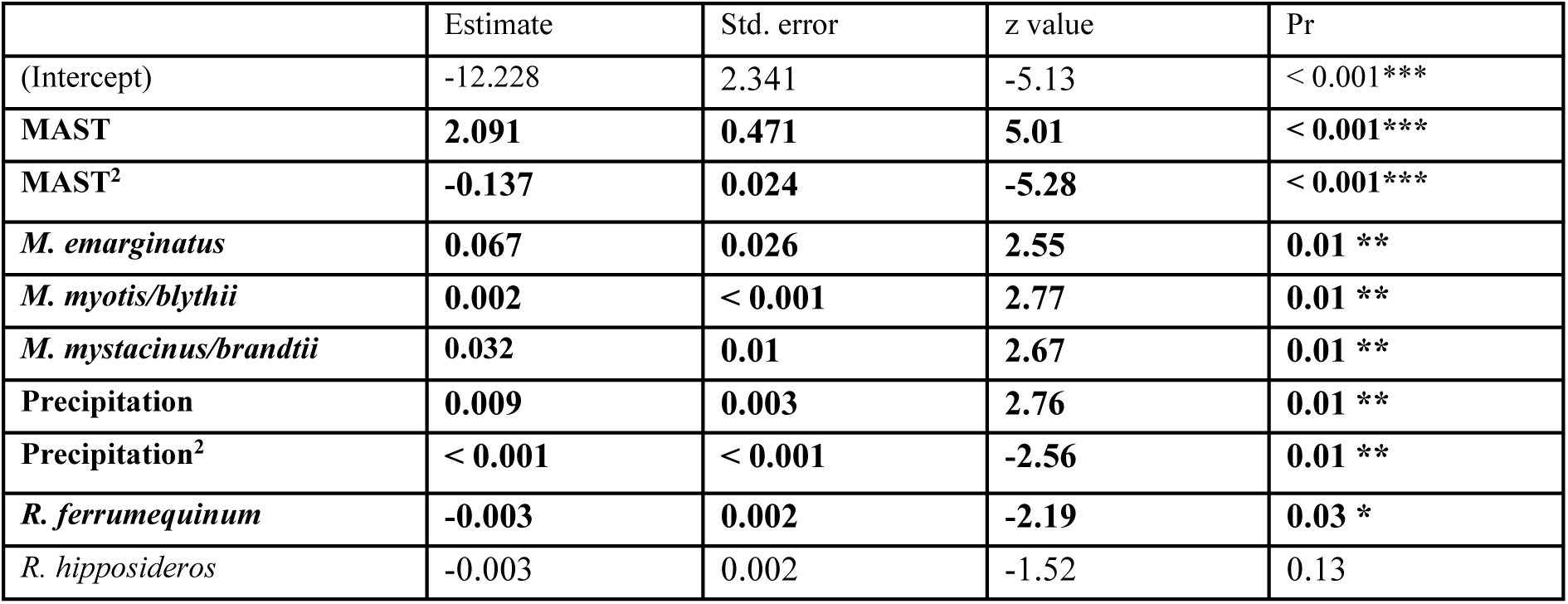
Results of the logistic regression model with lowest AICc (523.95) as selected by multi-model comparison. The response variable is whether signs of WND (fungal growth on the wings and/or muzzles) have ever been visually observed on a live bat at the site (1 = yes, 0 = no). The explanatory variables are species abundances in hibernation sites, as well as MAST and precipitation with their quadratic terms. Statistically significant variables (Wald Chi-square; p < 0.05) are bolded.

### 2.4 Extrapolation and transferability

As detailed above, our reference data set (i.e., the data we used for building the model) was collected in Europe and we applied this model beyond Europe. This means the model is being transferred to regions that may have different environmental conditions and/or potential bat host species, whose susceptibility to WND might not be known. Therefore, we first evaluated the transferability of our model theoretically, assessing geographic and host species transferability separately. We then validated our model empirically, simultaneously evaluating both geographic and host species transferability.

#### 2.4.1 Theoretical transferability

First, given that we applied our model to every cell of a raster map of the world, we checked that cells predicted to be WND-suitable harboured environmental conditions that were similar to those encountered in the recent (1970-2000) climatic conditions of the 448-point reference dataset geographically spread across Europe (see section ‘Data and data acquisition’). This was to ensure that the predictions of suitable cells are made within the range of the reference dataset. To achieve this, we calculated the multivariate environmental similarity surface (MESS) as described by Elith et al. (2010) using the mess function of the R dismo package (Hijmans et al. 2023). The MESS calculation represents how similar a point is to the reference dataset, with respect to the two environmental predictor variables herein used, MAST and precipitation. Negative values represent sites where at least one variable falls outside the range of environments observed in the reference set, indicating novel environments where predictions might not be accurate (Dormann et al. 2012). We calculated the percentage of cells with negative MESS values in relation to the predicted WND-suitability. To that end, we carried out the MESS analysis after masking the raster layers for WND-suitability equal or above a given threshold, the thresholds being considered starting at 0 until 1 by steps of 0.01. We performed these analyses for the recent (1970-2000) and future climatic conditions (2061-2080) separately.

Second, most of the host bat species that were included in our model have distributions that do not extend beyond the Palaearctic. Therefore, justifying the transfer of this model to regions lacking a subset or all of these species is necessary. As highlighted in the introduction, hibernation in contaminated hibernacula under favourable abiotic conditions appears to be the main risk factor rather than species-, genus- or family identity. This is for example illustrated by the fact that all European species for which a minimum of 10 individuals have been sampled have been diagnosed positive for *P. destructans* infection and the WND diagnostic histopathological lesions (e.g. Wibbelt et al. 2013, Zukal et al. 2014, 2016). Therefore, given the evidence gathered so far, it is most probable that most hibernating bat species are susceptible to some extent. We nevertheless mapped the worldwide distribution of several species-groups for which we have some information about WND susceptibility. We then calculated the percentage of areas with WND-suitability greater than 0.2 overlapping with the four different species groups investigated. The species-groups mapped were: 1) all species that have been recorded with WND (histopathological lesions confirmed), 2) all species that have been recorded with *P. destructans* infection, 3) all species in the bat genera included in the model (*Myotis, Rhinolophus, Plecotus, Barbastella*), 4) all species in the bat genus classically harbouring the highest *P. destructans* load and lesions (i.e. *Myotis*). Bat species distributions were recovered from the IUCN website as shape files (https://www.iucnredlist.org/; accessed 20.07.2023) and analysed in R.

#### 2.4.2 Empirical transferability

Next, we conducted two empirical validations to assess the transferability of our models to different regions and species concurrently. Utilising datasets from North America, we examined how well independent empirical datasets aligned with the predictions of our models. Notably, North America was chosen due to its geographic separation from Europe, where the model’s reference data originated. This was crucial because it enabled an external validation process that reflects real-world conditions more accurately than internal validations alone (e.g. cross-validations; Dormann et al. 2012). Moreover, given that there are no bat species common to both continents, this evaluation provided a robust test of the model’s generalizability.

First, we tested if the WND-related mortality data obtained from the North American WHISPers event sharing database (Wildlife Health Sharing Partnership 2023) were located in regions with higher WND-suitability than expected by chance. We used WND-related mortality rather than visual signs of the fungus as such a dataset does not exist for North America. Indeed, given that our model focuses on predicting optimal conditions for disease manifestation and severity, areas with the highest WND-suitability should correspond to those with the highest disease severity. This would translate to increased mortality in regions with susceptible species, as observed in North America following pathogen introduction. The difference in data used to train the model and test the model is a strength, as it allows for a more comprehensive evaluation of the model’s performance across different datasets and conditions. Additionally, this approach enhances the robustness of our analysis and provides valuable insights into the model’s applicability to real-world scenarios. To do so, we used data from the WHISPers database, the most comprehensive database available to date to centralise information on current and historic wildlife mortality and/or morbidity events in North America. On 30/11/2023, we queried the WHISPers database with ‘Event Type = Mortality/Morbidity’, ‘Event Diagnosis = White-Nose Syndrome’ and excluded one event (ID 15654) from late September 2008 as it is unlikely to represent Mortality/Morbidity related to WND. Five records out of 338 included two counties which were split (one data point per county), resulting in 343 Mortality/Morbidity events (293 in the USA, 50 in Canada). We excluded data on Canadian counties due to the unavailability of a raster layer needed for executing the analyses automatically (see below). Based on these data, we calculated the percentage of mortality events that were located in a county considered WND-suitable. A county was considered WND-suitable if it contained at least one cell with a WND-suitability value above a given threshold, the thresholds being considered starting at 0.01 until 0.99 by steps of 0.01. This procedure was carried out for the observed WHISPers dataset, but also with 1,000 simulated datasets acting as the null distribution. This null distribution corresponds to what would be expected if our model was no better than random at predicting WND-related Mortality/Morbidity. For the null distribution, we randomly selected the same number of counties (without replacement) as in the observed WHISPers dataset and conducted the analysis following the same procedures as with the observed dataset. Furthermore, we applied the identical procedure to predictions derived from data featuring a tenfold increase in the abundance of *M. emarginatus*, a species comparable in size to *M. lucifugus*, which exhibits the highest fungal load and mortality rates (Hoyt et al. 2021). This step was taken to ensure enhanced accuracy in mortality predictions by the model, particularly in a naïve population characterised by a higher prevalence of susceptible hosts, as observed in North America before the onset of WND (Frick et al. 2015).

Second, we adopted a comparative approach to assess the accuracy of our predictions. Given that our model focuses on predicting optimal conditions for disease manifestation and severity, we anticipated a better match between our model predictions and 1) WND-related Mortality/Morbidity, followed by 2) occurrence of the disease (as diagnosed by histologic lesions and *P. destructans* confirmation), and then 3) *P. destructans* presence alone. To test these predictions, we calculated the percentage of counties, meeting certain criteria (cf. below), that have at least one cell with a WND-suitability above a given threshold (thresholds considered started at 0.01 until 0.99 by steps of 0.01). This was achieved for counties 1) where *P. destructans* has been detected according to the ‘WNS spread map’ (=’*Pd* positive’; White Nose Syndrome Response team; queried on 30/11/2023), 2) where bats show characteristic histologic lesions and are *P. destructans* positive according to the ‘WNS spread map’ (=’Positive for WNS’; White Nose Syndrome Response team; queried on 30/11/2023), and, 3) where WND-related Mortality/Morbidity has been identified according to the WHISper database as detailed above for the first empirical validation.

## 3 RESULTS

### 3.1 Climatic preferences of European bat species during hibernation

Our dataset covered a wide range of climatic conditions (for MAST 0.5–17.9°C and for annual precipitation 299–1828 mm) in which the hibernation sites were located. The abundances of species/species groups within hibernation sites, as well as their distributions across the climatic conditions included in our dataset are described in Figure 3.

**Figure 3.**
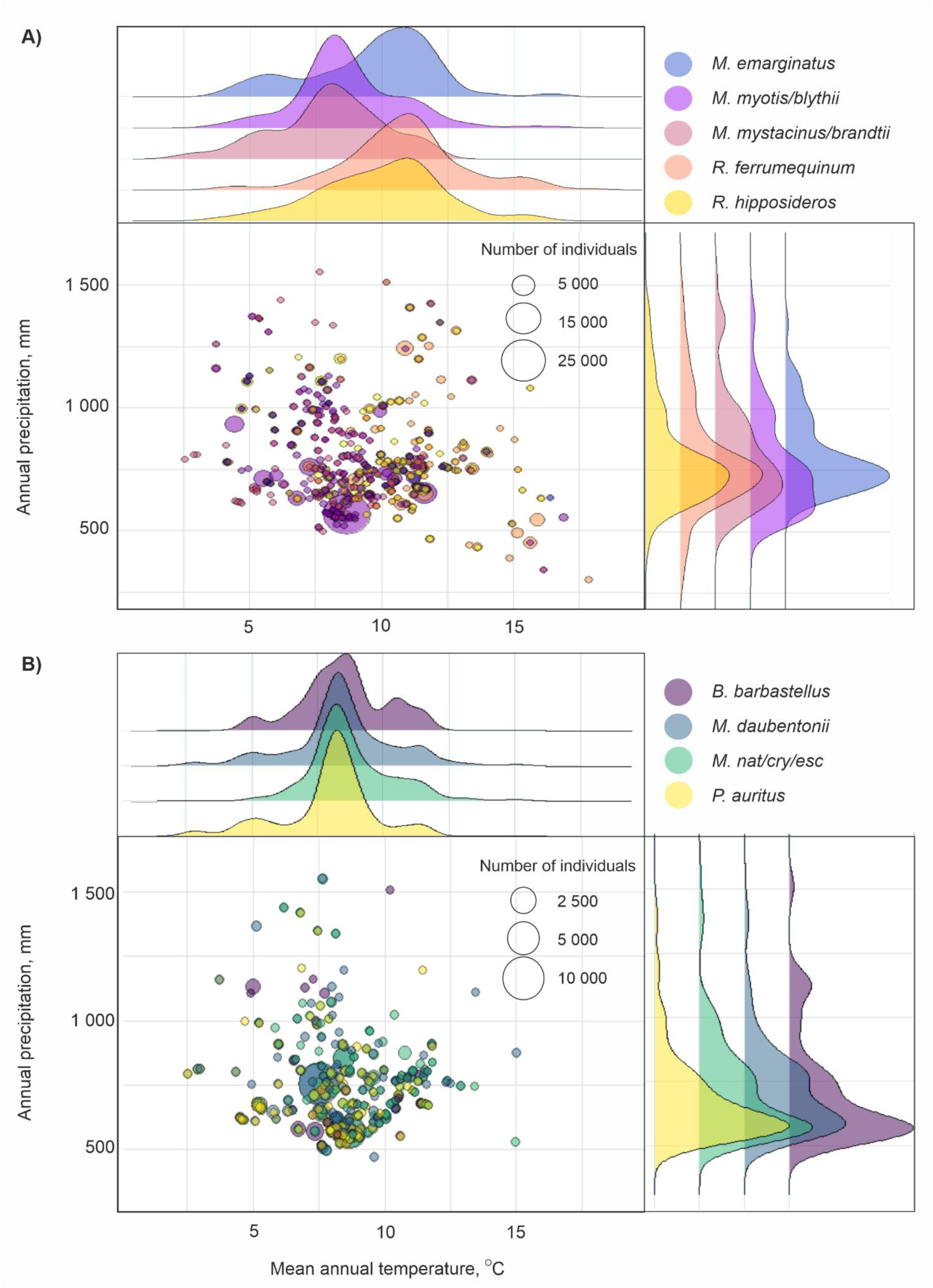
Bubble and density plots depicting the abundances of bat species/species groups hibernating across a range of climatic conditions. The size of the bubble is relative to the number of bats occupying a given hibernaculum. The density plots represent the distribution of hibernacula where the species is present. **A)** Distributions of five bat species included in the best model (see results): *M. emarginatus* (63 sites), *M. myotis/blythii* (262 sites, 60 148 individuals in total), *M. mystacinus/brandtii* (118 sites, 2 494 individuals in total), *R. ferrumequinum* (153 sites, 19 078 individuals in total), *R. hipposideros* (145 sites, 7 694 individuals in total). **B)** Distributions of four bat species not included in the best model: *Barbastella barbastellus* (53 sites, 4 737 individuals in total), *Myotis daubentonii* (221 sites, 35 604 individuals in total), *M. nattereri/crypticus/escalerai* (179 sites, 15 168 individuals in total), and *Plecotus auritus* (142 sites, 1 617 individuals in total).

### 3.2 Factors affecting the WND-suitability

Multi-model comparisons identified a logistic regression model with nine explanatory variables as the model with the lowest AICc (523.95; i.e., the best model). When comparing the full model, the best model, and three averaged models (averaged on models within < 2, < 4 and < 7 AICc of the best model), the estimated coefficients, their 85 % or 95 % confidence intervals, and significance were very similar, providing strong evidence that our inferences are robust to the model selection procedure (Tables S1-S8 Fig. S1). We therefore only present in the main text the results from the best model. Results from the best model revealed significant non-linear relationships between climatic conditions (MAST, precipitation) and WND-suitability (Figure 4A, B; Table 1). In addition, we found a positive significant association between the WND-suitability of hibernacula and the number of *Myotis emarginatus* (Table 1, Figure 4C), *M. myotis/blythii* (Table 1, Figure 4D), and *M. mystacinus/brandtii* (Table 1, Figure 4E), and a negative association with the number of *Rhinolophus ferrumequinum* (Table 1, Figure 4F). The number of *R. hipposideros*, selected in the best model, was negatively associated with the WND-suitability but was not significant (Wald Chi-square; Table 1).

**Figure 4.**
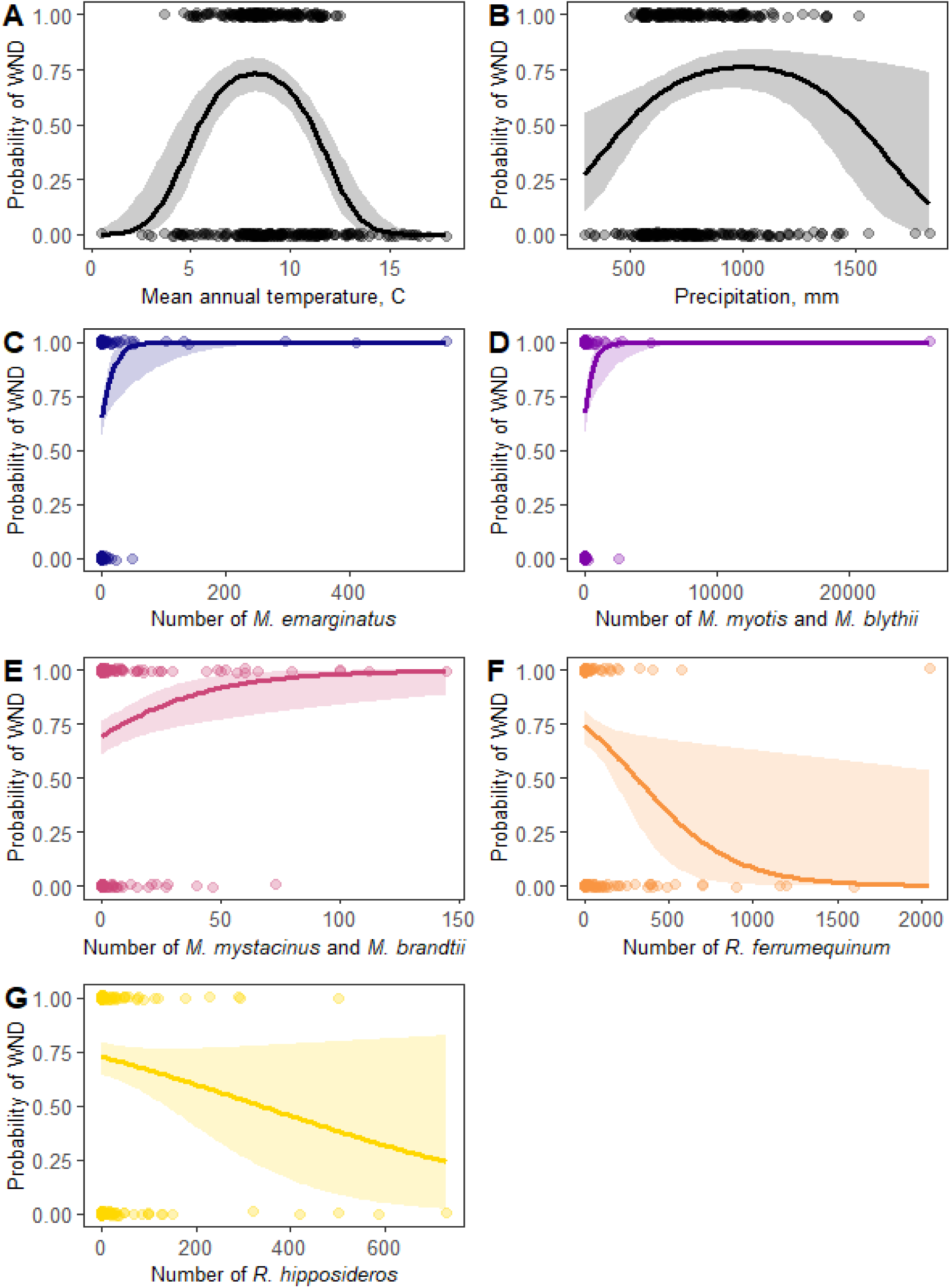
Effects of **A)** mean annual temperature, **B)** annual precipitation, and number of inhabiting **C)** *Myotis emarginatus*, **D)** *Myotis myotis/blythii*, **E)** *Myotis mystacinus/brandtii*, **F)** *Rhinolophus ferrumequinum*, and **G)** *Rhinolophus hipposideros* on the probability of observing visual signs of *P. destructans* growth (proxy for WND) on any live bat at a hibernation site. Coloured areas represent the 95% confidence intervals. Points represent the observed infection status in hibernacula (1= bats with visible *P. destructans*, 0 = no bats with visible *P. destructans*, jitter added for visual clarity).

### 3.3 Relationship between climatic conditions and environmental conditions within hibernation sites

We found a positive correlation between MAST and temperatures measured inside hibernacula (Pearson’s correlation coefficient r = 0.61, 95 % CI 0.54–0.67, p < 0.01, N = 356, Figure S5). According to the logistic regression analysis (Figure 3A), WND-suitability was highest in areas where MAST is 8.3°C. Fitting a linear model between hibernacula temperature and MAST revealed that MAST of 8.3°C equals 7.1°C inside hibernacula (95% CI 0.68—7.4, N= 356, Fig. S5). We also found a significant non-linear relationship between absolute humidity measured inside hibernation sites (p < 0.01, E = 8.177*10^-3^, SE = 3.061*10^-3^, t = 2.671) as well as its quadratic term (p = 0.03, t = -2.271, E = -3.689*10^-6^, SE=1.624*10^-6^). According to the model, the highest predicted absolute humidity (6.98 g/m^3^, 95 % CI = 6.982-7.895) occurred at 1051 mm annual precipitation (Fig. S6).

### 3.4 Predictions for WND-suitable areas now and in the future

Our model shows that under recent climatic conditions (1970-2000), 9.7 % of raster cells have a WND-suitability higher than 0.2. In addition to the current known distribution of WND in Eurasia and North America, climatic conditions may be suitable for the manifestation of the disease in several areas, including southern America, southern Australia, and New Zealand (Figure 5A). That being said, in terms of surface, the Northern hemisphere has 14 (recent) and 29 times (future) more area with WND-suitability > 0.2 than the southern hemisphere, highlighting the fact that the disease is primarily affecting the Northern hemisphere.

**Figure 5.**
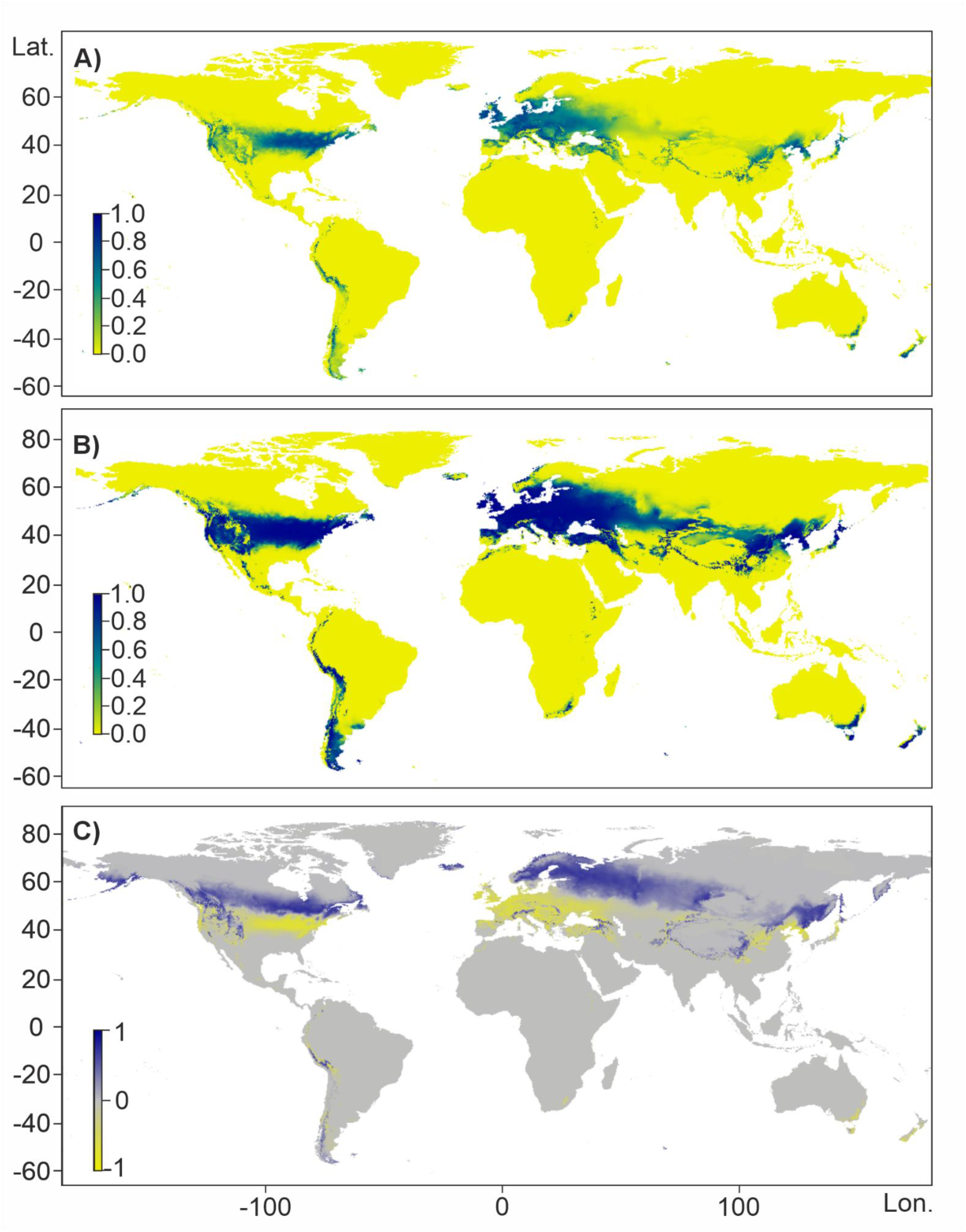
**A)** The global distribution of areas potentially at risk for WND in current climatic conditions, according to our model predictions computed using the average species abundance from Europe. **B)** The areas potentially at risk according to our model predictions computed by increasing the abundance of *M. emarginatus* 10-fold to demonstrate the effect of a high number of susceptible individuals. Other species are kept constant. **C)** Predicted shift in the areas suitable for WND globally during the years 2061–2080 under climate change model ACCESS-CM2, Shared Socio-economic Pathway 3-7.0., with the average species abundance from Europe. Yellow denotes areas where, compared to the present situation, the environmental suitability will decrease in the future while blue denotes areas where environmental suitability for WND will increase.

Multiplying the number of *M. emarginatus* 10-fold, while keeping the rest of the species composition constant, resulted in a considerable expansion in predicted WND-suitable areas (+171 %), illustrating the strong effect of local host abundance for the manifestation of disease (Figure 5B; Fig. S2, S7, S8). Besides extending the range, for suitable cells (i.e. with an average WND-suitability > 0.2), the average suitability increased from 0.49 to 0.68.

Applying our model to future climatic conditions showed a notable shift in the distribution of WND towards the north in the Palearctic and North America by the years 2061–2080 (Figure 5C). When compared to the recent climatic conditions, the number of raster cells with a WND-suitability higher than 0.2 will increase by 150 %, reaching 14.5 % of the world’s cells (Fig. S3). This increase in suitable range is the result of an imbalance between the number of cells with decreasing (6.8 %) and increasing suitability (11.5 %).

### 3.5 Validation of predictions

Transferability of our model was first assessed theoretically, considering geographic location and host species separately. For geographic transferability, the MESS analysis revealed an important similarity between environmental conditions of the reference dataset versus areas predicted to be WND-suitable, for both, recent and future conditions. This is illustrated by the fact that 97.5 % and 98.5 % of MESS values were positive for cells with WND-suitability > 0.2, and 100 % for WND-suitability > 0.27 (Fig. S9). So, all cells predicted to have a high WND-suitability have environmental conditions within the range of those encountered at the sites used for training the model. For host species transferability (Fig. S10-11), the distribution of all species that have been recorded with WND covers the large majority (recent: 89 %; future: 87 %) of areas predicted with WND-suitability > 0.2. This also applies to all species that have been recorded with *P. destructans* infection (recent: 93 %; future: 90 %). The distributions of some of the species recorded with *P. destructans* extend to the southern hemisphere, including the majority of the Andes in South America, where WND-suitability is elevated. When considering all species in the bat genera included in the model (*Myotis, Rhinolophus, Plecotus, Barbastella*), their distribution covered 95 % (recent) and 84 % (future) all of areas with WND-suitability > 0.2, which corresponds to all the main regions with high WND-suitability, except Tasmania and New Zealand. The distribution of the genus *Myotis* is less extensive than the four bat genera combined (*Myotis, Rhinolophus, Plecotus, Barbastella*) but is essentially similar in its coverage of regions with WND-suitability > 0.2 (recent: 93 %; future: 83 %).

Transferability of our model was then assessed empirically, dealing with geographic and host species concurrently using two different approaches. The first validation of our model and its transferability, comparing the accuracy of our predictions to US county-level mortality data (WHISPers event sharing database, Wildlife Health Sharing Partnership 2023) showed that our model systematically predicted the mortality status better than expected by chance (permutation test, p < 0.001), irrespective of the threshold used for the WND-suitability (Figure 6A). Increasing *M. emarginatus* to mimic a naïve population with a higher abundance of susceptible hosts, as observed in North America, further increased the accuracy with which our model predicted mortality in North America (Figure 6B).

**Figure 6.**
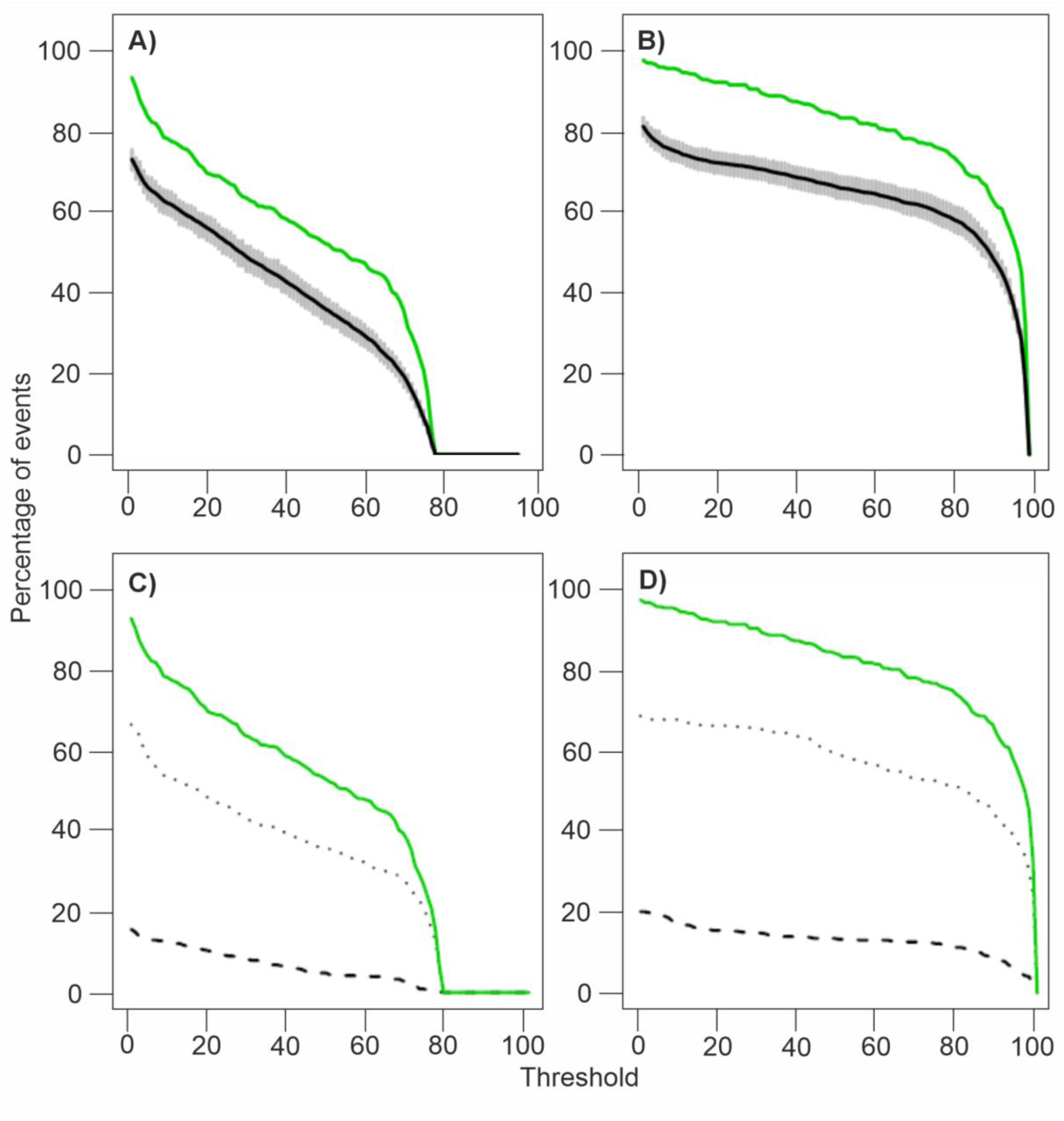
Validation of the predictions of our model. For each panel, the percentage of events is depicted for different thresholds of WND-suitability. WND-suitability is represented for the current climatic conditions for the average host species abundance (A and C) and for the average host species abundance bar the abundance of *M. emarginatus* which was multiplied by ten (B and D). In A and B, the green line represents the observed events of Mortality/Morbidity, the back line represents the average percentage of events under the null distribution (see text) with its 95 % highest density interval represented in grey. For C and D, the green lines are identical to A and B respectively, the dotted line represents events with the disease occurrence, and the dashed line represents events with the *P. destructans* presence.

As expected, irrespective of the threshold used, our second validation confirmed that predictions from our model are more related to Mortality/Morbidity than the presence of the disease alone, or simply presence of the pathogen (Figure 6C, D). This was true if we considered the average host-species abundances and even more so if we increased *M. emarginatus* to mimic a naïve population with a higher abundance of susceptible hosts (Figure 5 C versus D). Both of these validations emphasise the adequacy of our model, thereby confirming its transferability in terms of geography and host species to predict mortality from WND in naïve populations.

## 4 DISCUSSION

Our results underscore the importance of local climatic conditions and the presence, as well as abundance, of bat species in influencing the occurrence of WND in Europe. They also contribute further evidence that models developed and validated on one continent can effectively be applied to other continents, supported by the robust geographic and host species transferability observed in both theoretical and empirical assessments of our model. This offers new opportunities to better predict regions of the globe that could be conducive to specific diseases (Altizer et al. 2013), thereby aiding in conservation planning and disease prevention efforts, as recently achieved for *B. salamandrivorans*, a fungus causing chytridiomycosis in Amphibians (Fisher et al. 2020). Furthermore, because temperature is a critical factor influencing WND occurrence, predictions based on future climatic conditions (2061-2080) forecast drastic changes in the distribution of areas suitable for disease development. These potential shifts underscore the need for broad disease surveillance, ideally employing a minimally invasive monitoring method for bats, as suggested previously (Barlow et al. 2015, Barova et al. 2018, Fritze and Puechmaille 2018, Martínková et al. 2019, Hicks et al. 2020, Fritze et al. 2021b).

### 4.1 Environmental factors as key predictors of WND

With the largest dataset ever analysed for our purpose (448 sites), our study provides further clear evidence that a set of two environmental abiotic variables, MAST, and to a lesser extent precipitation, are important predictors of WND-suitability (i.e., the probability of *P. destructans* being visible on at least one bat at a site). Our model predicts that WND-suitability is highest in areas with a MAST of 8.3 °C, corresponding to 7.1 °C inside hibernation sites. Martínková et al. (Martínková et al. 2018) found maximal fungal infection temperatures between 3.8 and 7.8 °C, aligning with our results despite methodological and geographic differences. Their study covered a narrower temperature range (ca. 1–9 °C; N = 31 sites) compared to our broader dataset (0.5–17.9 °C; N = 448 sites) covering the full range of MAST used by hibernating bats. These differences, along with differences in species composition, could potentially explain why the maximum fungal load and amount of skin invasion predicted by Martínková et al. (Martínková et al. 2018) was lower (5–6 °C) than the 7.1 °C we found for the highest WND-suitability. Notwithstanding, the consistency between our current predictions and previous research on the occurrence of WND gives further credence to our approach and model. It confirms that there is a mismatch between the peak fungal growth temperature under laboratory conditions (12.5–15.8 °C) and the maximum growth/fungal load/disease severity observed in wild-living bats. Increasing temperatures lead to higher arousal frequencies in bats, which consequently increase grooming behaviour, thus interrupting fungal growth (Puechmaille et al. 2011b, Brownlee-Bouboulis and Reeder 2013, Horáček et al. 2014).

In addition to MAST, we found that annual precipitation also factored into the probability of WND, likely due to a causal link between precipitation and the humidity in hibernacula, the latter being an important environmental component for WND (Marroquin et al. 2017). Despite the potential difficulties in measuring humidity accurately, particularly as part of a large-scale citizen science project where sophisticated/accurate measuring devices are usually not available, and the fact that various other factors impacting humidity were not considered here (e.g., porosity and permeability of the geological surrounding, air flow, temporal variability, etc.) (Cigna 1968, Williams 2008). We found that the highest absolute humidity according to our model corresponded to 1051 mm annual precipitation. This is consistent with the precipitation determined optimal for WND occurrence according to our model (1000 mm annual precipitation). Our findings highlight that incorporating precipitation into predictive models for WND-suitability provides valuable information that cannot be solely captured by MAST. Due to the near ubiquitous variations in temperature (e.g. daily, seasonal, yearly, inter-annual) across ecosystems, it is usually challenging to identify the key climatic variables exerting the strongest pressure on a species (e.g. mean annual temperature, temperature seasonality, temperature of the coldest month, number of frost-free days). Considering a suite of variables, as often done with ecological niche modelling, is an option but disentangling the effect of the variable *per se* (e.g., temperature) is challenging as it is masked or blurred by *in situ* variations (e.g., within the day, day/night, seasonal, yearly) and collinearity with other variables. Working in underground sites, where conditions are more stable (Mammola et al. 2019), circumvents many of these challenges, enabling the development of models based on more causal relationships and mitigating several issues typically associated with correlative ecological niche models (Buckley et al. 2010, Kearney et al. 2010, Dormann et al. 2012, Johnson et al. 2019, Santini et al. 2021).

### 4.2 Effect of bat host species on the occurrence of WND in Europe

We found that WND-suitability was significantly increased by the abundance of three species/species groups in particular, *Myotis myotis/blythii*, *M. mystacinus/brandtii*, and *M. emarginatus*, reflecting the importance of available host composition for the disease to manifest. A positive association can be expected between WND-suitability and a species abundance if the species is susceptible to the disease. Fungal load explains most of the differences in the probability of observing visible *P. destructans* on bats. Therefore, all other things being equal, the probability of observing WND at a site should be related to the abundance of the most susceptible host species.

We observed a significant increase in WND-suitability associated with the abundance of three species or species groups: *Myotis myotis/blythii*, *M. mystacinus/brandtii*, and *M. emarginatus*. While all of these species have been previously recorded with WND, the species most often associated with WND in Europe is *M. myotis* (Martínková et al. 2010, Puechmaille et al. 2011b, Zukal et al. 2014, 2016). In fact, visible signs of WND appear to be rare in regions where *M. myotis/blythii* are rare or absent, for instance Great Britain, Ireland and southern Scandinavia (Puechmaille et al. 2011b, Ågren et al. 2012, Barlow et al. 2015). The lack of visual signs of WND in these areas appears to indeed be due to *M. myotis/blythii* not being present because the model predicts that the environmental conditions for the manifestation of the disease are met in these regions (Figure 4A). This underscores the potential role of *M. myotis/blythii* as pivotal contributors to the proliferation of *P. destructans*, thus playing a significant role in driving the local occurrence of WND in Europe. Amongst species investigated in Europe, *M. myotis/blythii* exhibit the highest fungal loads (Zukal et al. 2014). Given that fungal load largely explains variations in the probability of observing visible *P. destructans* on bats (Janicki et al. 2015), the association between WND-suitability and *M. myotis/blythii* abundance seems causal. In other words, *M. myotis/blythii* are sentinel species for the detection of WND. Furthermore, *M. myotis/blythii* contribute to the proliferation of *P. destructans* at hibernacula by replenishing the environmental reservoir at the end of the hibernation season (Puechmaille et al. 2011b, Hoyt et al. 2020, Fischer et al. 2020, 2022).

In addition to *M. myotis*/*blythii*, we uncovered two other species/species groups, *M. emarginatus* and *M. mystacinus/brandtii*, to have a positive association between their number in hibernacula and the probability of WND. Given that we found no significant multicollinearity between the three, their associations with WND are unlikely to be due to these species selecting the same hibernation sites. Currently, information on the relationship between WND and *M. brandtii* and *M. mystacinus* are limited and, given that our dataset did not allow to separate these two species (that may differ in their responses and/or disposition to infection), more studies are needed to determine the reasons behind the positive association our analysis revealed. Interestingly, most hibernacula occupied by *M. emarginatus* appear to have temperatures a few degrees above the optimal temperature for maximal fungal load/disease presence (Martínková et al. 2018; our study). We propose that, instead of environmental conditions, the positive relationship between *M. emarginatus* abundance and the occurrence of WND could be mostly due to its unique hibernation behaviour: the species is known to extend its hibernation period to as late as the middle of May (Dietz et al. 2009), allowing *P. destructans* to have more time for growth and propagation. This interpretation, which deserves further investigation, is corroborated by the fact that most individual *M. emarginatus* captured in June at one hibernation site in Italy still carried viable *P. destructans* (Garzoli et al. 2019, 2021). Besides, in some regions, *M. emarginatus* is the second most common species after *M. myotis* to show visible signs of *P. destructans* growth (Horáček et al. 2014). Hibernation phenology may also provide an explanation for why we did not find any association between the occurrence of WND and *M. nattereri* species complex. The hibernation period of *M. nattereri* is considerably short, taking place approximately from December/January to the beginning of March (Meier et al. 2022, Razgour 2023), possibly limiting *P. destructans* growth on the species. With regards to *M. daubentonii*, we identified no association with WND, despite the species having an extended hibernation period, typically spanning from September-October until March, in environmental conditions conducive to the proliferation of *P. destructans* (Meier et al. 2022). Our findings are consistent with data on fungal load, indicating that *M. daubentonii* has one of the lowest fungal loads among European *Myotis* species (Zukal et al. 2016). Notably, *M. daubentonii* is the only bat species for which infection by *Pd*-1, the *P. destructans* lineage accounting for approximately 95% of infections across host species, has not been documented (Fischer et al. 2025). Accordingly, the absence of a significant association between *M. daubentonii* and WND-suitability in our analyses is biologically coherent and further demonstrate that *M. daubentonii* has likely evolved alternative mechanisms to limit *Pd*-1 growth, and further studies would be required to elucidate their nature.

In contrast, we found a negative correlation between WND-suitability and the abundance of *Rhinolophus ferrumequinum*. The negative relationship between *R. ferrumequinum* and WND is in line with results from previous studies that have observed lower infection rates and fungal prevalence in the species in Asia and Europe (Hoyt et al. 2020). This has led to the suggestion of *Rhinolophus sp.* being more resistant towards infection by *P. destructans* than other bat species, or that it may be protected by microbes with antifungal properties (Li et al. 2022, Leng et al. 2026). In addition, our data indicates that environmental conditions in hibernation sites occupied by *R. ferrumequinum* were situated in climates with MAST above 10 °C (Figure 3), i.e., above 8.1 °C, the temperature with maximal probability for WND occurrence. In these conditions, bats have increased arousal frequencies compared to colder conditions (Ransome 1971, Park et al. 2003, Hope and Jones 2012), allowing them to control the fungus via grooming (Puechmaille et al. 2011b, Brownlee-Bouboulis and Reeder 2013). In addition, unlike many other bat species, including several *Myotis* spp., *Rhinolophus* species do not typically crawl on hibernaculum walls, thereby limiting contact with wall surfaces that serve as environmental reservoirs of spores (Fischer et al. 2022). This likely results in a lower initial infectious dose, which may consequently constrain the capacity of *P. destructans* to establish, proliferate, and develop within the host.

The fact that bat species are positively, negatively or not associated with WND means that changes in species composition alone are expected to affect the probability of WND occurrence (Laggan et al. 2023). All other things being equal, a site with large number of hibernating *M. emarginatus*, *M. myotis/blythii,* or *M. mystacinus/brandtii* will inevitably have a higher probability of WND presence than a site with large numbers of *R. ferrumequinum, R. hipposideros, M. daubentonii, M. nattereri* species complex, or *Barbastella barbastellus*. Presently, species interactions were not incorporated into our modelling framework, as our primary aim was to concentrate on the strongest effects acting at large scales.

### 4.3 Transferability of the model

Predicting the presence of species or diseases beyond their known range is crucial in various scenarios, particularly for anticipating the impact of new pathogen introductions to different regions or continents, and assessing the potential effects of climate change. Classically, these predictions pose challenges for two primary reasons, which can interact with each other. First, predictor variables from the range we aim to predict often partially fall outside the range of values in the reference dataset, necessitating extrapolation. Second, if the model relies on predictor variables that are merely correlated with the response variable rather than causally linked, these correlations may vary across different locations, possibly resulting in inaccurate extrapolations (Dormann et al. 2012). Although there is no ground truth to which we can compare our model predictions, our different approaches to evaluate transferability, theoretical and empirical, converged to validate our projections. We intentionally selected a small number of abiotic predictor variables with causal links to the response variable and results from the multivariate environmental similarity surface confirmed that all areas predicted with WND-suitability > 0.27 were within the range of values observed in the reference dataset, hence without extrapolation. This was true for both, recent and future climatic conditions. For regions with low WND-suitability, especially when MAST > 18–20 °C, beyond values observed in the reference dataset, the model extrapolated. Nevertheless, the extrapolation is supported by solid biological evidence. Laboratory experiments have demonstrated that the pathogen does not grow above 20 °C (Verant et al. 2012), and observations indicate that bats do not exhibit extended torpor use above such temperatures (Webb et al. 1996), hence we are confident in the predictions that disease suitability is very low in such conditions.

Regarding theoretical host species transferability, over 87 % of the area predicted with WNS-suitability > 0.2 overlaps with one or several species that have already been diagnosed with WND. If we instead consider species that have been infected, or species that belong to a genera included in our model, or species that belong to the single genus most commonly affected (*Myotis*), we are systematically above 83 %, for recent or future climatic conditions. These distributional data of potential hosts strongly support the idea that the distribution of the fungus causing WND, which is neither species-, genus- nor family-specific (Zukal et al. 2014, Hoyt et al. 2021, Wu et al. 2025), is unlikely to be geographically limited by bat hosts at large scales. That being said, while our study focused on estimating effect sizes for bat species within Europe, future research could employ the same modelling framework to compare these estimates across species on different regions where the fungus is present (i.e. Asia, North America). Once accomplished, models incorporating the true local species composition and abundance could be developed beyond Europe to yield more precise information concerning the potential impact of WND on local bat populations.

The solid results from the evaluation of theoretical transferability are in agreement with the empirical validation we conducted, evaluating simultaneously geographic and host species transferability from Europe to North America. This provides further evidence that applying the European trained model to predict WND-suitability in other continents with non-overlapping bat species performs well. Indeed, tests on empirical data provided evidence that predictions made from a model exclusively trained from European data predicted mortality from the disease in North America well (Figure 5). The difference in data utilised for model development (visual detection of *P. destructans* on live bats in Europe, where mortality rarely occurs even when the disease is severe) and assessment (WND-related mortality in North America, where severe disease may cause mortality in susceptible species) is a strength, as it enables a robust evaluation of the model’s performance across varied datasets and circumstances. The probability of observing visible *P. destructans* on bats is strongly related to fungal load (Janicki et al. 2015, Fritze et al. 2021b), and wing damage, a measure of disease severity (Fritze et al. 2021a, 2021b). Therefore, it is expected that in areas where the disease causes mortality (e.g. North America), the probability of observing visible *P. destructans* on bats is also a good proxy for mortality. This expectation is supported by our geographical extrapolation of the model to North America. Further support comes from the second empirical evaluation of the model, showing that in fact, our model predictions for North America better match the observed WND-related mortality, rather than the simple presence of the disease (with or without associated mortality; Figure 6C-D). Therefore, it is important to keep in mind that our model specifically predicts WND-suitability as a proxy of disease severity as measured by dermatological damage (Fritze et al. 2021b) and not only the presence/absence of the disease itself. Notwithstanding, given that different bat host species with comparable dermatological damage can have different disease outcome (e.g. Pikula et al. 2017, Whiting-Fawcett et al. 2021, 2025), our model does not predict the disease outcome per se (e.g. death, recovery, etc.).

### 4.4 Predictions for WND-suitability through space and time

Predictions over the world emphasise that the Northern hemisphere, particularly Europe, North America and to a lesser extent parts of Asia have the largest areas with high WND-suitability. In fact, WND-suitability is 14 and 29 times more widespread in the Northern hemisphere compared to the Southern hemisphere, for recent and future climatic conditions respectively. That being said, the majority of bat species in the Northern hemisphere have by now been exposed to *P. destructans*, the causative agent of WND. This is not the case in the Southern hemisphere where the pathogen is thought to be absent. Hence, if introduced, it could significantly affect local bat populations as recently evidenced in North America. Therefore, our model gives further credence to the concern of *P. destructans* introduction to the southern parts of South America, Southern Australia, and New Zealand (Turbill and Welbergen 2019, Lilley et al. 2020), and highlights the need for proper hygiene when visiting hibernation sites (Zhelyazkova et al. 2020, 2025) especially in areas that are climatically suitable for the manifestation of the disease. Our predictions also indicated areas with high WND-suitability over parts of Lesotho and South Africa where several species in the *Rhinolophus*, *Miniopterus* and *Myotis* genera are known to use extended torpor (Monadjem et al. 2020).

Regarding Asia, regions with high WND-suitability are restricted to a narrow crescent going from Central Asia, through the Northern part of Southern Asia, into Eastern Asia with the largest suitable areas being in the Northern and Northeastern parts of China. The latter corresponds to the region where *P. destructans* was first confirmed in Asia, and where visible signs of WND have also been observed (Hoyt et al. 2016, 2021). Regrettably, there is limited published data regarding the occurrence of WND in Asia. Apart from eleven sites in Russia where coordinates are provided along with histopathological information (Zukal et al. 2016, Zahradníková et al. 2018, Kovacova et al. 2018), we were not able to recover from the published literature further data associating site coordinates with WND status. This lack of information makes it challenging to test our model using data from Asia in a similar way we did for North America. That being said, our predictions encompass several extensive regions where *P. destructans* has been detected such as Northern and Northeastern parts of China, Honshu in Japan, the Far East in Russia, or the Korean Peninsula (Kovacova et al. 2018, Hoyt et al. 2021, Kim et al. 2022). Our predictions in Asia and elsewhere should be considered as conservative in the sense that our model does not predict the most mild forms of the disease (i.e. limited to moderate dermatological damage), as evidenced with the data transferability analysis for North America.

The presence and abundance of a few key species can affect the probability to detect visible signs of the fungus on live bats (Janicki et al. 2015) and this is mirrored by the fact that mimicking high abundance of a key species increased WND-suitability (Figure 5). A higher abundance of hosts means that more pathogen propagules will be released in the environmental reservoir, increasing the infectiousness of the site (Fischer et al. 2022, Laggan et al. 2023, Vanderwolf et al. 2025). A higher infectiousness results in more individuals being infected by a higher dose, leading to more pronounced disease (Frick et al. 2017). Therefore, host abundance, especially those that are most susceptible to the pathogen, and disease severity are causally linked. This causal link, captured by the model, therefore predicts that when the abundance of some key species increases, WND-suitability increases. This is particularly true when we multiplied the number of *M. emarginatus* 10-fold to illustrate how the abundance of susceptible hosts affects the predictions, as would be expected in a naïve population that comes into contact with the pathogen for the first time. This 10-fold change was picked to mimic the population sizes present in North America before the onset of the WND (Frick et al. 2015). Taken together, our model can be used to reliably predict mortality events in naïve populations, even when species are non-overlapping such as between Europe and North America as demonstrated herein. This underscores the practicality of our model in identifying regions warranting caution to prevent the introduction of the pathogen.

Interestingly, our predicted distribution of the disease in North America, computed using the average species abundances in Europe, does not spread out as far in the south, where the fungus has been recorded. We propose that this phenomenon might be linked to the high abundance of susceptible hosts, especially *M. lucifugus* and *M. septentrionalis*, which enabled the efficient proliferation of the fungus (Laggan et al. 2023). This occurred despite the environmental conditions not being optimally conducive for the disease. This is supported by the significant expansion and improved accuracy in our predicted range of WND in North America, resulting from multiplying the abundance of *M. emarginatus*, the species with the strongest effect on WND presence in Europe (Figure 5). These results also illustrate that the presence and abundance of one species can have a significant impact on WND-suitability, and therefore, highlight the importance of host composition in WND dynamics. Our findings reveal an interesting system in which to test the Eltonian noise hypothesis, suggesting that coarse-scaled abiotic factors like temperature and precipitation play a significant role in determining distributions at broad scales. Meanwhile, ecological interactions and species’ effects on resources are influential in shaping distributions at finer spatial scales (Soberón and Nakamura 2009).

Applying our model to a climate change scenario resulted in dramatic shifts in the potential distribution range of WND by the years 2061–2080, showing that climate change has the potential of putting new bat populations at risk of WND in a relatively short time. Although the future climate is predicted to be warmer at the global scale, the geographical area of occurrence of the disease caused by the psychrophilic *P. destructans* is expected to be higher (Figure 4C). This can be attributed largely to the expanding availability of suitable areas in the Northern Hemisphere. While our results imply that WND has the potential to heavily affect areas in Northern Europe, WND may not be able to manifest unless the distribution range of the most important host species also shifts/expands north. These results also highlight the need for improvements to gain an even deeper insight into the complex host-pathogen-environment interactions that shape the disease (Whiting-Fawcett et al. 2025), and lays the premise to instigate such further studies to provide a holistic view of the interactions shaping temperate bat communities.

## 5 CONCLUSIONS

No previous study has undertaken the task to model the global distribution of WND. Therefore, our model represents a powerful initial approximation and working hypothesis for identifying both current and future areas suitable for the disease. The model identifies regions where environmental conditions favour disease outbreak, provided both host and pathogen are present. Our findings emphasise the critical role of two key environmental factors, mean annual surface temperature and precipitation, as governing WND manifestation across large scales, mirroring patterns observed in other fungal wildlife diseases like chytridiomycosis (Rohr and Raffel 2010, Murray et al. 2011). Furthermore, our model aligns with previous chytridiomycosis predictions (Rohr and Raffel 2010) in suggesting that climate change may significantly reshape and expand suitable areas for WND. This focus on environmental conditions is crucial because we demonstrated that the vast diversity of bat species globally means that susceptible populations are likely present wherever these conditions become favourable. Therefore, the model offers a broader perspective for our general understanding of the disease and prioritising conservation efforts. The model can guide targeted surveillance efforts in high-risk regions, allowing for preventative measures to be implemented and the introduction of the pathogen into new areas to be prevented. Additionally, the model can be used within an adaptive management framework to strategically collect future data and refine its accuracy over time. By providing a foundation for future research and guiding critical conservation strategies, this WND distribution model holds significant promise for mitigating the devastating impacts of this disease.

