## Supplementary information for "Climatic factors and host species composition at hibernation sites drive the incidence of bat fungal disease"

### SUPPLEMENTARY MATERIAL 1:

#### Additional information on model selection and comparison, model validation, environmental measurements and model predictions

**Table S1.** Results of the logistic regression model including all explanatory variables. The response variable is information on whether signs of WND (fungal growth on the wings and/or muzzles) have ever been visually observed on a live bat at the site (1 = yes, 0 = no). The explanatory variables are species abundances in hibernation sites, as well as MAST and precipitation with their quadratic terms. Statistically significant variables ( p < 0.05) are bolded.

| **Coefficients:** | **Estimate** | **Std.Error** | **zvalue** | **Pr(>\|z\|)** | **Significance level** |
|---|---|---|---|---|---|
| (Intercept) | -12.330 | 2.4230 | -5.088 | < 0.001 | *** |
| ***M. myotis/blythii*** | **0.002** | **0.001** | **2.773** | **< 0.01** | ** |
| *M. daubentonii* | < 0.001 | 0.001 | 0.366 | 0.71 | |
| *M. nat/esc/cry* | 0.002 | 0.003 | 0.664 | 0.51 | |
| *P. auritus* | 0.036 | 0.034 | 1.065 | 0.29 | |
| ***R. ferrumequinum*** | **-0.003** | **0.002** | **-2.168** | **0.03** | * |
| *R. hipposideros* | -0.003 | 0.002 | -1.510 | 0.13 | |
| ***M. emarginatus*** | **0.071** | **0.027** | **2.633** | **< 0.01** | ** |
| *M. mystacinus/brandtii* | 0.023 | 0.014 | 1.677 | 0.09 | . |
| *B. barbastellus* | -0.002 | 0.003 | -0.648 | 0.52 | |
| **Precipitation** | **0.009** | **0.003** | **2.899** | **< 0.01** | ** |
| **Precipitation$^2$** | **< 0.001** | **< 0.001** | **-2.669** | **< 0.01** | ** |
| **MAST** | **2.013** | **0.416** | **4.839** | **< 0.001** | *** |
| **MAST$^2$** | **-0.122** | **0.024** | **-5.109** | **< 0.001** | *** |

**Table S2.** Results of the logistic regression model selected by multi-model comparison of AICc values. The response variable is information on whether signs of WND (fungal growth on the wings and/or muzzles) have ever been visually observed on a live bat at the site (1 = yes, 0 = no). The explanatory variables are species abundances in hibernation sites, as well as MAST and precipitation with their quadratic terms. Statistically significant variables (p < 0.05) are bolded.

| | Estimate | Std. error | z value | Pr | Significance level |
|---|---|---|---|---|---|
| (Intercept) | -12.228 | 2.341 | -5.13 | < 0.001 | *** |

| | | | | | |
|---|---|---|---|---|---|
| **MAST** | **2.091** | **0.471** | **5.01** | **< 0.001** | *** |
| **MAST²** | **-0.137** | **0.024** | **-5.28** | **< 0.001** | *** |
| ***M. emarginatus*** | **0.067** | **0.026** | **2.55** | **0.01** | * |
| ***M. myotis/blythii*** | **0.002** | **< 0.001** | **2.77** | **< 0.01** | ** |
| ***M. mystacinus/brandtii*** | **0.032** | **0.01** | **2.67** | **< 0.01** | ** |
| **Precipitation** | **0.009** | **0.003** | **2.76** | **< 0.01** | ** |
| **Precipitation²** | **< 0.001** | **< 0.001** | **-2.56** | **<0.01** | * |
| ***R. ferrumequinum*** | **-0.003** | **0.002** | **-2.19** | **0.03** | * |
| *R. hipposideros* | -0.003 | 0.002 | -1.52 | 0.13 | |

**Table S3.** Results of the averaged logistic regression analysis gained from models within 2 AICc from the model with the lowest AICc.

| | Estimate | Std. Error | Adjusted SE | z value | Pr(>\|z\|) | Significance level |
|---|---|---|---|---|---|---|
| (Intercept) | -12.300 | 2.412 | 2.418 | 5.086 | < 0.001 | *** |
| **MAST** | **2.072** | **0.417** | **0.418** | **4.952** | **< 0.001** | *** |
| **MAST²** | **-0.125** | **0.024** | **0.024** | **5.207** | **< 0.001** | *** |
| ***M. emarginatus*** | **0.067** | **0.026** | **0.026** | **2.514** | **0.012** | * |
| ***M. myotis/blythii*** | **0.002** | **0.001** | **0.001** | **2.774** | **0.006** | ** |
| ***M. mystacinus/brandtii*** | **0.030** | **0.013** | **0.013** | **2.353** | **0.019** | * |
| **Precipitation** | **0.009** | **0.003** | **0.003** | **2.753** | **0.006** | ** |
| **Precipitation²** | **0.000** | **0.000** | **0.000** | **2.555** | **0.011** | * |
| ***R. ferrumequinum*** | **-0.004** | **0.002** | **0.002** | **2.212** | **0.027** | * |
| *R. hipposideros* | -0.002 | 0.002 | 0.002 | 0.906 | 0.365 | |
| *P. auritus* | 0.010 | 0.024 | 0.024 | 0.416 | 0.677 | |
| *M. nat/esc/cry* | 0.001 | 0.002 | 0.002 | 0.329 | 0.742 | |
| *M. daubentonii* | 0.000 | 0.001 | 0.001 | 0.225 | 0.822 | |
| *B. barbastellus* | 0.000 | 0.001 | 0.001 | 0.163 | 0.870 | |

**Table S4**. Results of the averaged logistic regression analysis gained from models within 4 AICc from the model with the lowest AICc.

| | Estimate | Std. Error | Adjusted SE | z value | Pr(>\|z\|) | |
|---|---|---|---|---|---|---|
| (Intercept) | -12.270 | 2.413 | 2.420 | 5.069 | 0.000 | *** |
| **MAST** | **2.055** | **0.418** | **0.419** | **4.909** | **0.000** | *** |
| **$MAST^2$** | **-0.124** | **0.024** | **0.024** | **5.169** | **0.000** | *** |
| ***M. emarginatus*** | **0.067** | **0.027** | **0.027** | **2.526** | **0.012** | * |
| ***M. myotis/blythii*** | **0.002** | **0.001** | **0.001** | **2.775** | **0.006** | ** |
| ***M. mystacinus/brandtii*** | **0.028** | **0.014** | **0.014** | **1.998** | **0.046** | * |
| **Precipitation** | **0.009** | **0.003** | **0.003** | **2.767** | **0.006** | ** |
| **$Precipitation^2$** | **0.000** | **0.000** | **0.000** | **2.567** | **0.010** | * |
| ***R. ferrumequinum*** | **-0.004** | **0.002** | **0.002** | **2.213** | **0.027** | * |
| *R. hipposideros* | -0.002 | 0.002 | 0.002 | 0.876 | 0.381 | |
| *P. auritus* | 0.015 | 0.029 | 0.029 | 0.535 | 0.593 | |
| *M. nat/esc/cry* | 0.001 | 0.002 | 0.002 | 0.360 | 0.719 | |
| *M. daubentonii* | 0.000 | 0.001 | 0.001 | 0.273 | 0.785 | |
| *B. barbastellus* | 0.000 | 0.002 | 0.002 | 0.297 | 0.766 | |

**Table S6.** Results of the averaged logistic regression analysis gained from models within 7 AICc from the model with the lowest AICc.

| | Estimate | Std. Error | Adjusted SE | z value | Pr(>\|z\|) | |
|---|---|---|---|---|---|---|
| (Intercept) | -12.000 | 2.558 | 2.564 | 4.681 | **< 0.001** | *** |
| **MAST** | **2.029** | **0.419** | **0.420** | **4.831** | **< 0.001** | *** |
| **MAST$^2$** | **-0.123** | **0.024** | **0.024** | **5.092** | **< 0.001** | *** |
| ***M. emarginatus*** | **0.068** | **0.027** | **0.027** | **2.537** | **0.011** | * |
| ***M. myotis/blythii*** | **0.002** | **0.001** | **0.001** | **2.726** | **0.006** | ** |
| ***M. mystacinus/brandtii*** | **0.026** | **0.016** | **0.016** | **1.653** | **0.098** | . |
| **Precipitation** | **0.009** | **0.004** | **0.004** | **2.336** | **0.020** | * |
| **Precipitation$^2$** | **0.000** | **0.000** | **0.000** | **2.198** | **0.028** | * |
| ***R. ferrumequinum*** | **-0.003** | **0.002** | **0.002** | **2.096** | **0.036** | * |
| *R. hipposideros* | -0.002 | 0.002 | 0.002 | 0.841 | 0.400 | |
| *P. auritus* | 0.018 | 0.031 | 0.031 | 0.593 | 0.553 | |
| *M. nat/esc/cry* | 0.001 | 0.002 | 0.002 | 0.377 | 0.706 | |
| *M. daubentonii* | 0.000 | 0.001 | 0.001 | 0.292 | 0.770 | |
| *B. barbastellus* | -0.001 | 0.002 | 0.002 | 0.319 | 0.750 | |

**Table S7.** Key for component models

| Code | Term |
|---|---|
| 1 | *B. barbastellus* |
| 2 | MAST |
| 3 | MAST$^2$ |
| 4 | *M. daubentonii* |
| 5 | *M. emarginatus* |
| 6 | *M. myotis/blythii* |
| 7 | *M mystacinus/brandtii* |
| 8 | *M. nattereri/* |
| 9 | *P. auritus* |
| 10 | Precipitation |

| | |
|---|---|
| 11 | Precipitation$^2$ |
| 12 | *R. ferrumequinum* |
| 13 | *R. hipposideros* |

**Table S8.** Models within 2, 4, and 7 AICc from the model with the lowest AICc value.

| **Component model** | **df** | **logLik** | **AICc** | **delta** | **weight** |
|---|---|---|---|---|---|
| 2+3+5+6+7+10+11+12+13 | 10 | -251.72 | 523.95 | 0 | 0.08 |
| 2+3+5+6+7+10+11+12 | 9 | -253.13 | 524.67 | 0.71 | 0.06 |
| 2+3+5+6+7+9+10+11+12+13 | 11 | -251.03 | 524.67 | 0.72 | 0.06 |
| 2+3+5+6+7+8+10+11+12+13 | 11 | -251.14 | 524.88 | 0.92 | 0.05 |
| 2+3+4+5+6+7+10+11+12+13 | 11 | -251.27 | 525.14 | 1.19 | 0.04 |
| 2+3+5+6+7+9+10+11+12 | 10 | -252.46 | 525.43 | 1.47 | 0.04 |
| 1+2+3+5+6+7+10+11+12+13 | 11 | -251.43 | 525.47 | 1.52 | 0.04 |
| 2+3+5+6+7+8+10+11+12 | 10 | -252.49 | 525.48 | 1.53 | 0.04 |
| 2+3+5+6+7+8+9+10+11+12+13 | 12 | -250.49 | 525.69 | 1.74 | 0.03 |
| 2+3+4+5+6+7+10+11+12 | 10 | -252.66 | 525.82 | 1.86 | 0.03 |
| 2+3+4+5+6+7+9+10+11+12+13 | 12 | -250.63 | 525.98 | 2.02 | 0.03 |
| 1+2+3+5+6+7+10+11+12 | 10 | -252.86 | 526.22 | 2.27 | 0.03 |
| 1+2+3+5+6+7+9+10+11+12+13 | 12 | -250.77 | 526.26 | 2.31 | 0.03 |
| 2+3+5+6+7+8+9+10+11+12 | 11 | -251.87 | 526.34 | 2.39 | 0.02 |
| 1+2+3+5+6+7+8+10+11+12+13 | 12 | -250.86 | 526.43 | 2.47 | 0.02 |
| 1+2+3+4+5+6+7+10+11+12+13 | 12 | -250.98 | 526.67 | 2.72 | 0.02 |
| 2+3+4+5+6+7+9+10+11+12 | 11 | -252.04 | 526.68 | 2.73 | 0.02 |
| 2+3+4+5+6+7+8+10+11+12+13 | 12 | -251 | 526.71 | 2.76 | 0.02 |
| 2+3+4+5+6+9+10+11+12+13 | 11 | -252.21 | 527.02 | 3.07 | 0.02 |
| 1+2+3+5+6+7+9+10+11+12 | 11 | -252.22 | 527.05 | 3.1 | 0.02 |
| 1+2+3+5+6+7+8+10+11+12 | 11 | -252.23 | 527.07 | 3.11 | 0.02 |
| 1+2+3+5+6+7+8+9+10+11+12+13 | 13 | -250.24 | 527.31 | 3.36 | 0.01 |
| 2+3+4+5+6+7+8+10+11+12 | 11 | -252.35 | 527.32 | 3.36 | 0.01 |
| 1+2+3+4+5+6+7+10+11+12 | 11 | -252.39 | 527.38 | 3.43 | 0.01 |

| | | | | | |
|---|---|---|---|---|---|
| 2+3+4+5+6+7+8+9+10+11+12+13 | 13 | -250.32 | 527.47 | 3.52 | 0.01 |
| 2+3+5+6+9+10+11+12+13 | 10 | -253.5 | 527.5 | 3.55 | 0.01 |
| 1+2+3+4+5+6+7+9+10+11+12+13 | 13 | -250.37 | 527.57 | 3.62 | 0.01 |
| 1+2+3+5+6+7+8+9+10+11+12 | 12 | -251.64 | 527.99 | 4.04 | 0.01 |
| 2+3+4+5+6+9+10+11+12 | 10 | -253.81 | 528.12 | 4.16 | 0.01 |
| 2+3+4+5+6+7+8+9+10+11+12 | 12 | -251.7 | 528.12 | 4.17 | 0.01 |
| 2+3+5+6+8+9+10+11+12+13 | 11 | -252.81 | 528.23 | 4.28 | 0.01 |
| 1+2+3+4+5+6+7+8+10+11+12+13 | 13 | -250.71 | 528.27 | 4.31 | 0.01 |
| 1+2+3+4+5+6+7+9+10+11+12 | 12 | -251.8 | 528.31 | 4.36 | 0.01 |
| 1+2+3+4+5+6+9+10+11+12+13 | 12 | -251.88 | 528.49 | 4.53 | 0.01 |
| 2+3+4+5+6+8+9+10+11+12+13 | 12 | -251.93 | 528.58 | 4.62 | 0.01 |
| 2+3+5+6+9+10+11+12 | 9 | -255.18 | 528.77 | 4.81 | 0.01 |
| 1+2+3+4+5+6+7+8+10+11+12 | 12 | -252.09 | 528.9 | 4.95 | 0.01 |
| 1+2+3+5+6+9+10+11+12+13 | 11 | -253.16 | 528.92 | 4.97 | 0.01 |
| 1+2+3+4+5+6+7+8+9+10+11+12+13 | 14 | -250.06 | 529.1 | 5.14 | 0.01 |
| 2+3+5+6+7+12 | 7 | -257.43 | 529.11 | 5.16 | 0.01 |
| 2+3+5+6+7+12+13 | 8 | -256.42 | 529.17 | 5.22 | 0.01 |
| 2+3+5+6+8+9+10+11+12 | 10 | -254.43 | 529.36 | 5.41 | 0.01 |
| 2+3+5+6+7+10+11+13 | 9 | -255.52 | 529.46 | 5.51 | 0.01 |
| 2+3+4+5+6+10+11+12+13 | 10 | -254.52 | 529.55 | 5.6 | 0 |
| 2+3+4+5+6+8+9+10+11+12 | 11 | -253.51 | 529.62 | 5.67 | 0 |
| 1+2+3+4+5+6+9+10+11+12 | 11 | -253.51 | 529.62 | 5.67 | 0 |
| 1+2+3+5+6+8+9+10+11+12+13 | 12 | -252.49 | 529.7 | 5.75 | 0 |
| 1+2+3+4+5+6+7+8+9+10+11+12 | 13 | -251.47 | 529.78 | 5.83 | 0 |
| 2+3+5+6+7+10+12+13 | 9 | -255.71 | 529.83 | 5.88 | 0 |
| 1+2+3+4+5+6+8+9+10+11+12+13 | 13 | -251.61 | 530.07 | 6.11 | 0 |
| 2+3+5+6+7+10+12 | 8 | -256.87 | 530.07 | 6.11 | 0 |
| 2+3+5+6+7+9+10+11+13 | 10 | -254.8 | 530.11 | 6.15 | 0 |
| 2+3+5+6+7+8+10+11+13 | 10 | -254.84 | 530.17 | 6.22 | 0 |
| 1+2+3+5+6+9+10+11+12 | 10 | -254.86 | 530.23 | 6.28 | 0 |

| | | | | | |
|---|---|---|---|---|---|
| 2+3+5+6+7+8+12 | 8 | -256.96 | 530.25 | 6.3 | 0 |
| 2+3+4+5+6+7+12 | 8 | -257 | 530.33 | 6.38 | 0 |
| 2+3+5+6+7+8+12+13 | 9 | -256 | 530.4 | 6.45 | 0 |
| 1+2+3+5+6+7+12 | 8 | -257.05 | 530.42 | 6.47 | 0 |
| 1+2+3+5+6+7+12+13 | 9 | -256.01 | 530.43 | 6.48 | 0 |
| 2+3+4+5+6+7+12+13 | 9 | -256.01 | 530.44 | 6.49 | 0 |
| 2+3+4+5+6+10+11+12 | 9 | -256.04 | 530.49 | 6.54 | 0 |
| 2+3+4+5+6+7+10+11+13 | 10 | -255.05 | 530.61 | 6.66 | 0 |
| 2+3+5+6+7+11+12+13 | 9 | -256.2 | 530.82 | 6.87 | 0 |
| 2+3+5+6+7+10+11 | 8 | -257.27 | 530.87 | 6.92 | 0 |
| 1+2+3+5+6+8+9+10+11+12 | 11 | -254.13 | 530.87 | 6.92 | 0 |
| 2+3+5+6+7+11+12 | 8 | -257.28 | 530.89 | 6.94 | 0 |
| 2+3+5+6+7+9+12 | 8 | -257.3 | 530.92 | 6.97 | 0 |
| 1+2+3+4+5+6+10+11+12+13 | 11 | -254.16 | 530.92 | 6.97 | 0 |
| 2+3+5+6+7+8+9+10+11+13 | 11 | -254.17 | 530.95 | 6.99 | 0 |

**Table S9.** Estimates of regression coefficients of the full model when using all 448 data points (Full dataset) or jackknifed datasets (448 datasets with 447 data points each). For the jackknifed estimates, the average across jackknifed datasets is presented along with its standard deviation (sd). The difference between the full dataset estimate and the average of the estimates for the jackknifed datasets is also presented (‘Difference’).

| | Jackknifed dataset | | Full dataset | |
|---|---|---|---|---|
| coef | Estimate | sd | Estimate | Difference |
| (Intercept) | -1.23E+01 | 1.14E-01 | -1.23E+01 | 1.71E-03 |
| Mmyobly | 1.97E-03 | 6.26E-04 | 1.94E-03 | 2.61E-05 |
| Mdau | 2.80E-04 | 1.53E-05 | 2.80E-04 | 3.33E-07 |
| Mnatsl | 2.10E-03 | 2.53E-04 | 2.09E-03 | 8.27E-06 |
| Paur | 3.61E-02 | 1.51E-03 | 3.61E-02 | 2.21E-05 |
| Rfer | -3.45E-03 | 2.04E-04 | -3.45E-03 | 6.72E-06 |
| Rhip | -2.98E-03 | 8.98E-05 | -2.98E-03 | 1.89E-07 |
| Mema | 7.06E-02 | 2.63E-03 | 7.06E-02 | 1.92E-05 |

| | | | | |
|---|---|---|---|---|
| Mmysbra | 2.33E-02 | 7.27E-04 | 2.33E-02 | 2.12E-07 |
| Bbar | -1.87E-03 | 2.42E-03 | -1.76E-03 | 1.07E-04 |
| PREC | 9.36E-03 | 1.55E-04 | 9.35E-03 | 2.64E-06 |
| PREC^2 | -4.60E-06 | 8.25E-08 | -4.60E-06 | 1.27E-09 |
| MAT | 2.01E+00 | 2.20E-02 | 2.01E+00 | 4.13E-05 |
| MAT^2 | -1.22E-01 | 1.25E-03 | -1.22E-01 | 1.21E-06 |

**Table S10.** Correlation between predictions for the recent climatic conditions (1970-2000; below the diagonal) or the future conditions (2061-2080; above the diagonal) for the full model, the best model, and three averaged models (coefficient averaged for models < 2, < 4 and < 7 AICc of the best model). Correlation coefficients were rounded to the nearest fifth digit.

| | Full | Best | AICc < 2 | AICc < 4 | AICc < 7 |
|---|---|---|---|---|---|
| Full | - | 0.99923 | 0.99947 | 0.99964 | 0.9994 |
| Best | 0.99918 | - | 0.99996 | 0.99999 | 0.9998 |
| AICc < 2 | 0.99944 | 0.99996 | - | 0.99998 | 0.99988 |
| AICc < 4 | 0.99961 | 0.9999 | 0.99998 | - | 0.9999 |
| AICc < 7 | 0.99929 | 0.99979 | 0.99986 | 0.99987 | - |

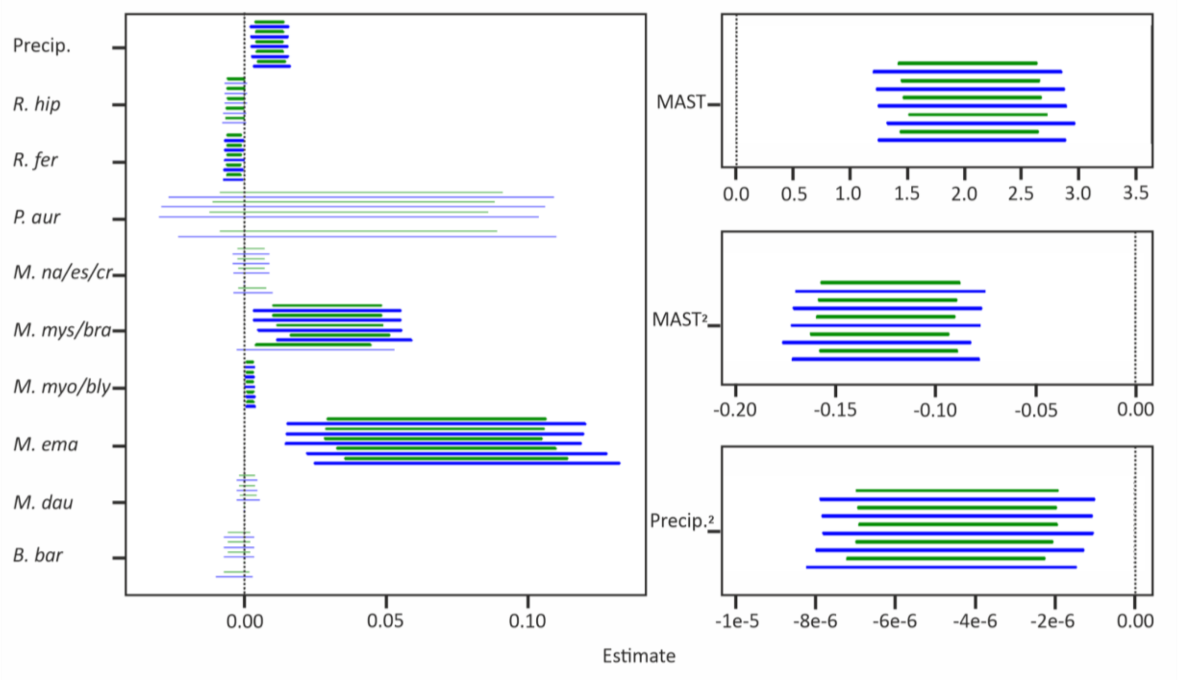


**Figure S1**. The coefficient plot illustrates estimates-specific confidence intervals (CIs) for each variable (depicted on the Y-axis) and each model (see below). Horizontal lines in green and blue represent 85% and 95% CIs, respectively. For each variable, the CIs, arranged from bottom to top, correspond to CIs of estimates from the full model, the best model (with the lowest AICc), and model-averaged models within 2, 4, and 7 AICc of the best model. Thick and thin lines distinguish CIs that overlap or do not overlap with zero. Variables are grouped according to the magnitudes of their estimates, resulting in varying scales along the X-axis for the different panels. In instances where a variable is omitted from a model (*P.aur*, *M.na/es/cr*, *M.dau* & *B.bar*, in the best model) or when the confidence interval cannot be computed (M.dau in the full model), a vacant space is visible.

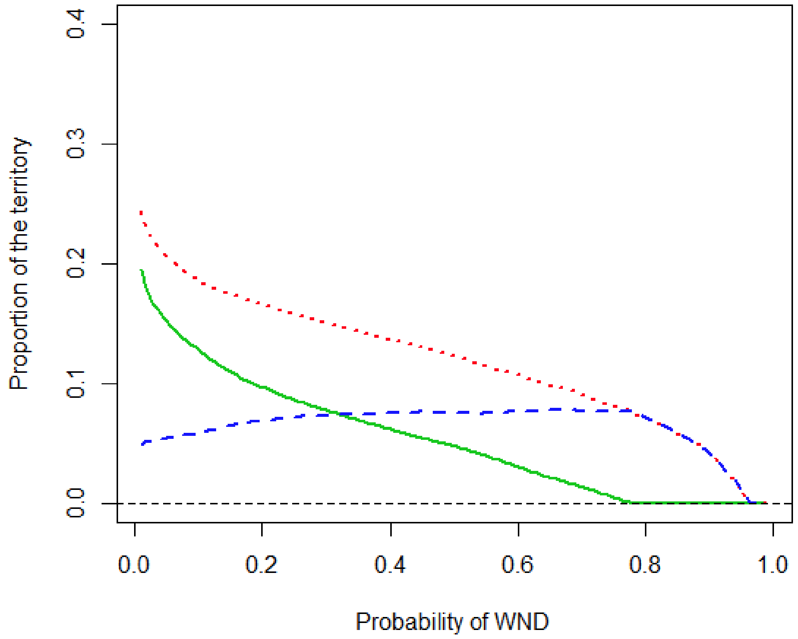


**Figure S2.** Proportion of the land cells in relation to the probability of WND presence (=WND-suitability). The green line represents the recent climatic conditions (1970-2000), the red dotted line represents the recent climatic conditions with the number of *M. emarginatus* multiplied by ten, and the blue dashed line represents the difference between the recent climatic conditions with the observed versus increased (x10) number of *M. emarginatus*.

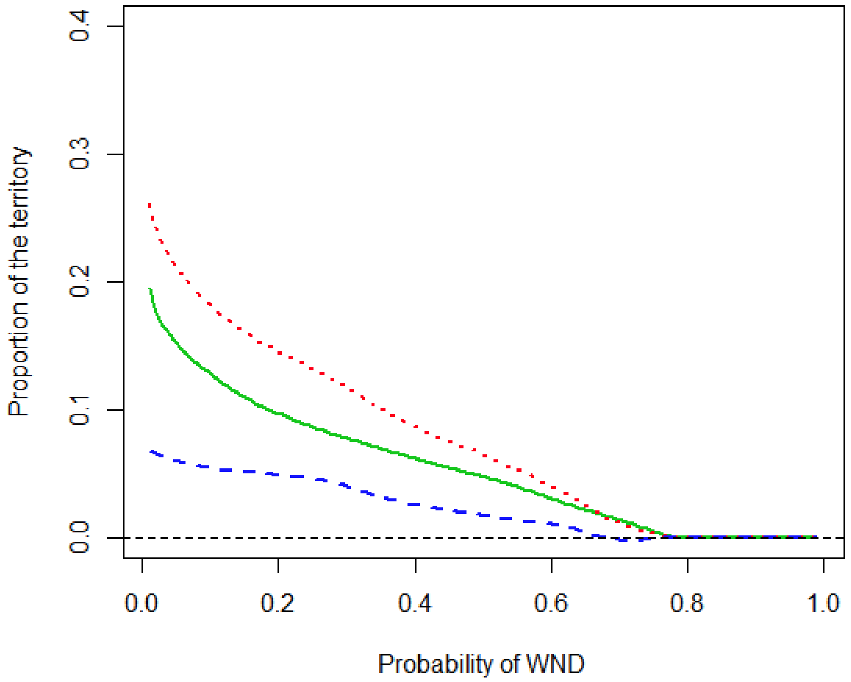


**Figure S3**. Proportion of the land cells in relation to the probability of WND presence (=WND-suitability). The green line represents the recent climatic conditions (1970-2000), the red dotted line represents the future climatic conditions (2061–2080) and the blue dashed line represents the difference between the future and the recent climatic conditions.

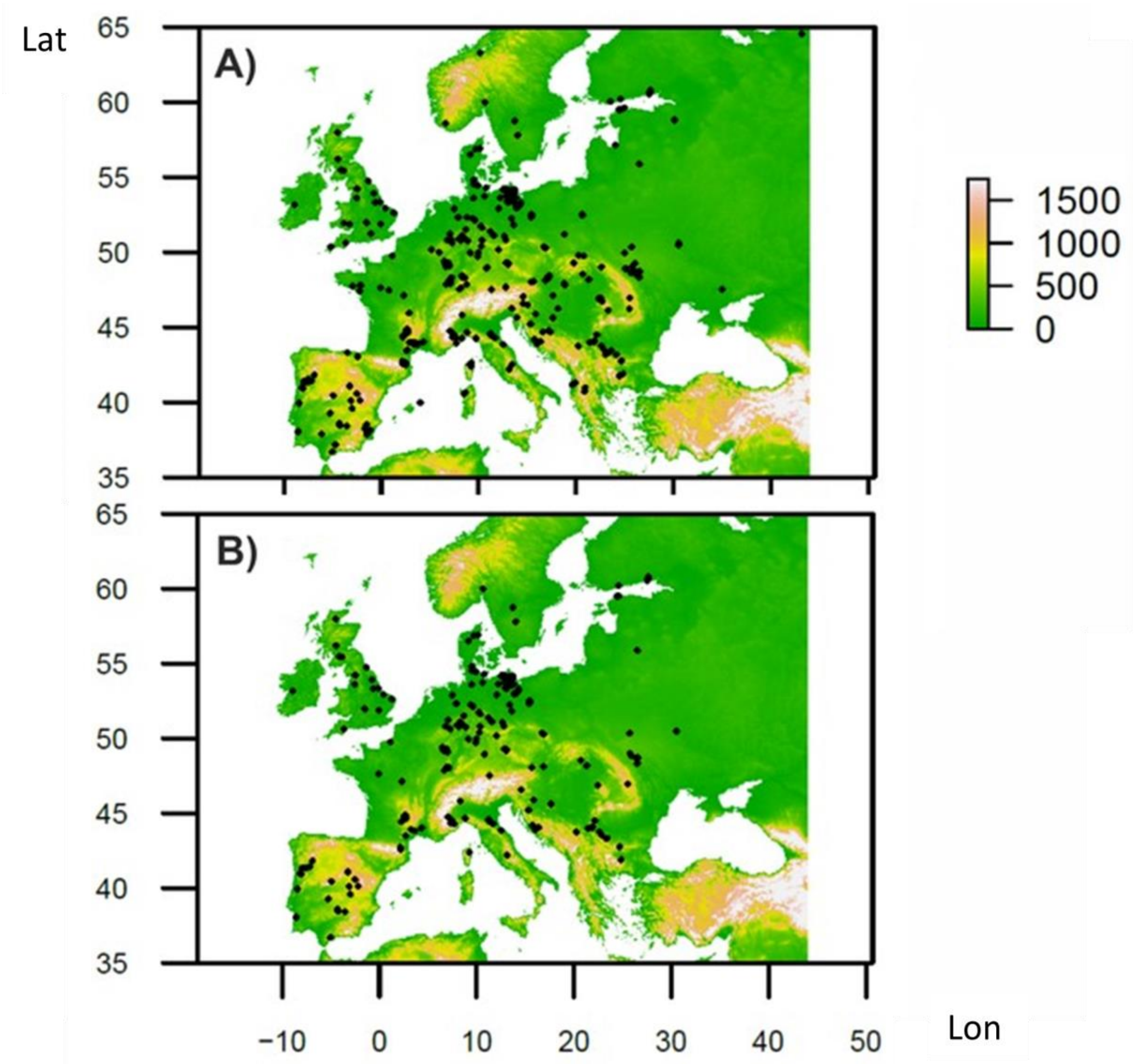


**Figure S4.** Map of sites where environmental measures for **A)** temperature inside hibernacula and **B)** relative humidity inside hibernacula.

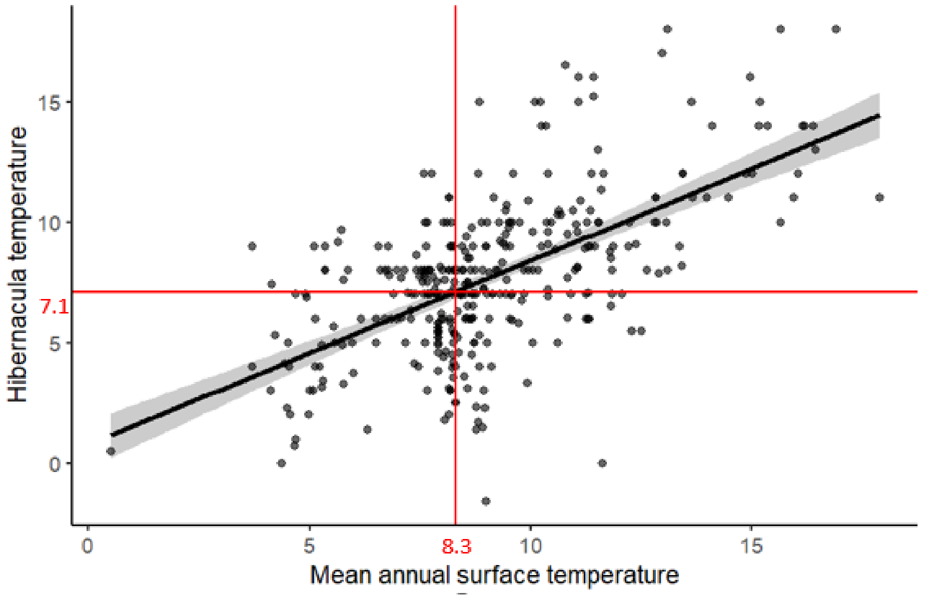


**Figure S5.** The relationship between the mean annual surface temperature and temperature measured inside hibernacula (Spearman's correlation coefficient rho = 0.54, 95% CI = 0.47–0.61, $p < 0.01$, N = 356 sites). The mean annual surface temperature of 8.3°C, determined optimal for the occurrence of WND in our study, corresponds to 7.1°C (95% CI=6.8–7.4) inside hibernacula according to the fitted linear regression model.

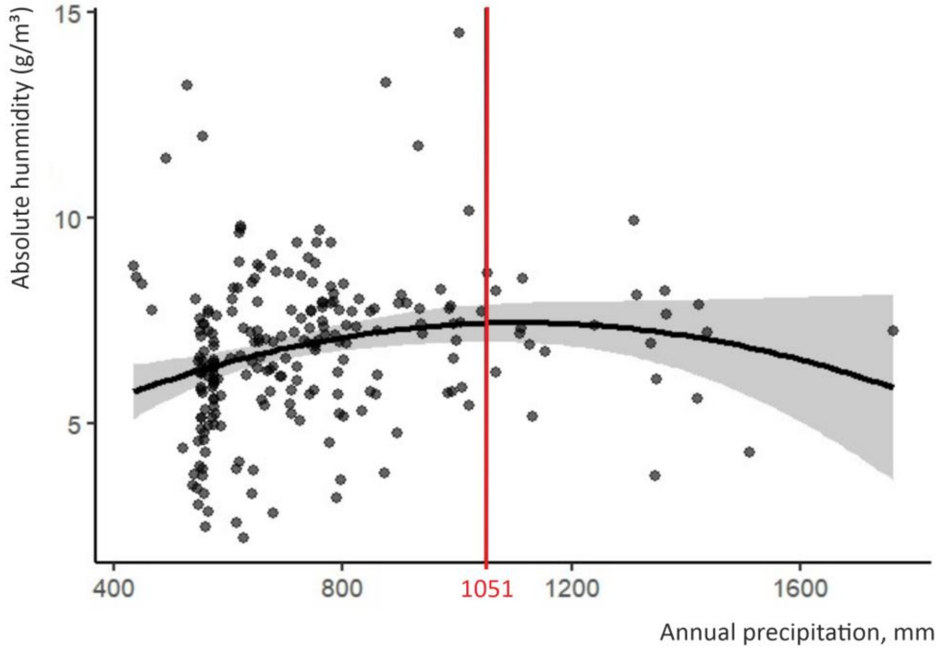


**Figure S6.** To investigate the relationship between annual precipitation and absolute humidity within hibernation sites, we fit a generalized linear model with absolute humidity (N=228 sites) as the response variable, and precipitation and its quadratic term as explanatory variables. We found a non-linear relationship between absolute humidity and precipitation ($p < 0.01$, $E = 8.177*10^{-3}$, $SE=3.061*10^{-3}$, $t=2.671$) and its quadratic term ($p = 0.03$, $t= -2.271$, $E = -3.689*10^{-6}$, $SE=1.624*10^{-6}$). The highest predicted absolute humidity (6.98 g/m$^3$, 95 % CI=6.982-7.895) occurred at 1051 mm annual precipitation.

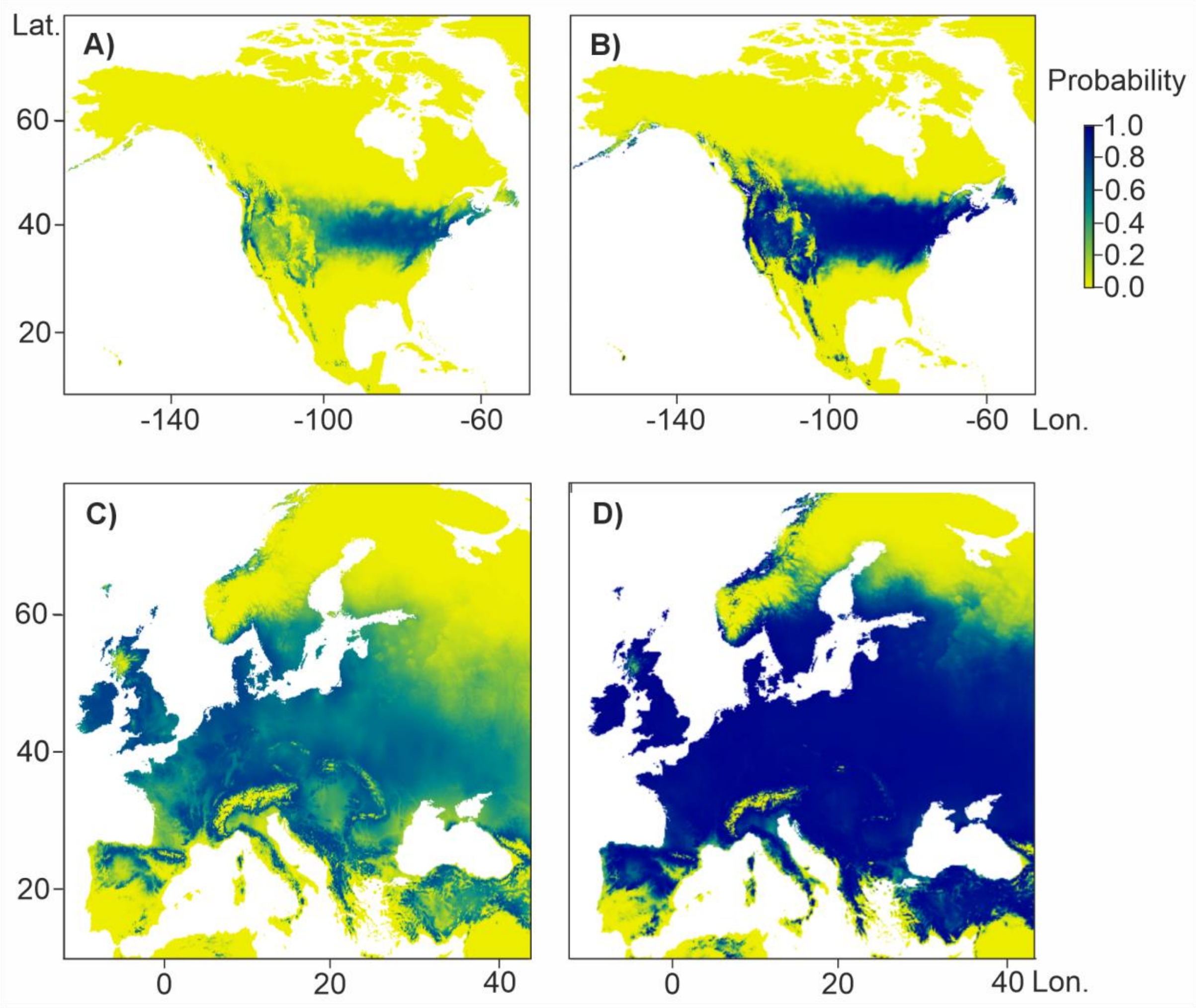


**Figure S7. A)** Predicted probability of WND occurrence (=WND-suibability) in North America computed using the average European species composition. **B)** Predicted probability of the occurrence of WND in North America computed with a 10-fold increase in *Myotis emarginatus* to demonstrate the effect of the abundance of susceptible hosts to disease distribution. **C)** Predicted probability of WND occurrence in Europe computed using the average European species composition. **D)** Predicted probability of the occurrence of WND in Europe computed with a 10-fold increase in *M. emarginatus* to demonstrate the effect of the abundance of susceptible hosts on WND-suibability.

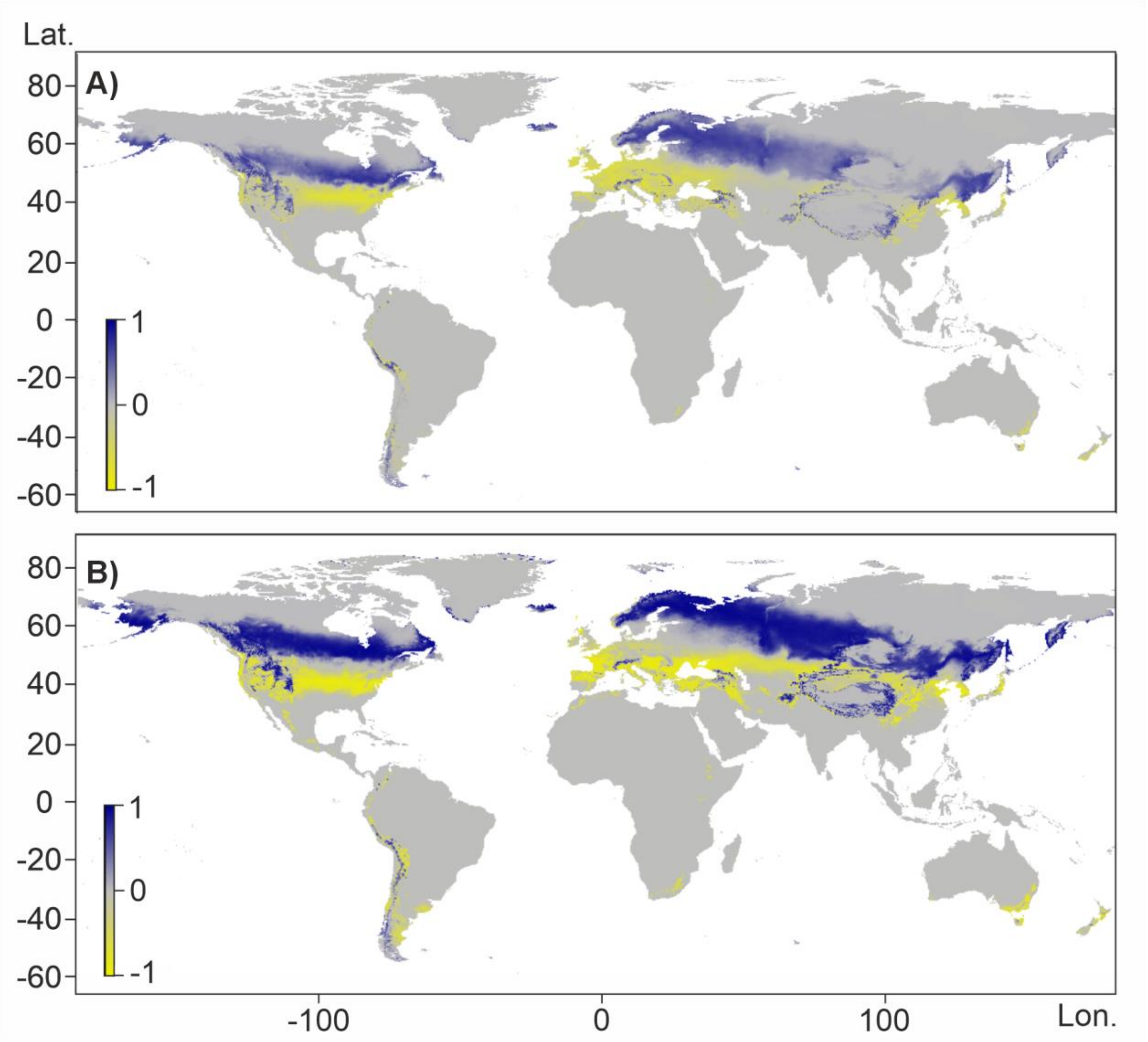


**Figure S8.** Difference between the predictions of the current and future distribution of WND computed with **A)** average European species abundance and **B)** a 10-fold increase in *M. emarginatus* to demonstrate the effect of the abundance of susceptible hosts on WND-suitability. Yellow denotes areas where, compared to the present situation, the environmental suitability will decrease in the future while blue denotes areas where environmental suitability for WND will increase.

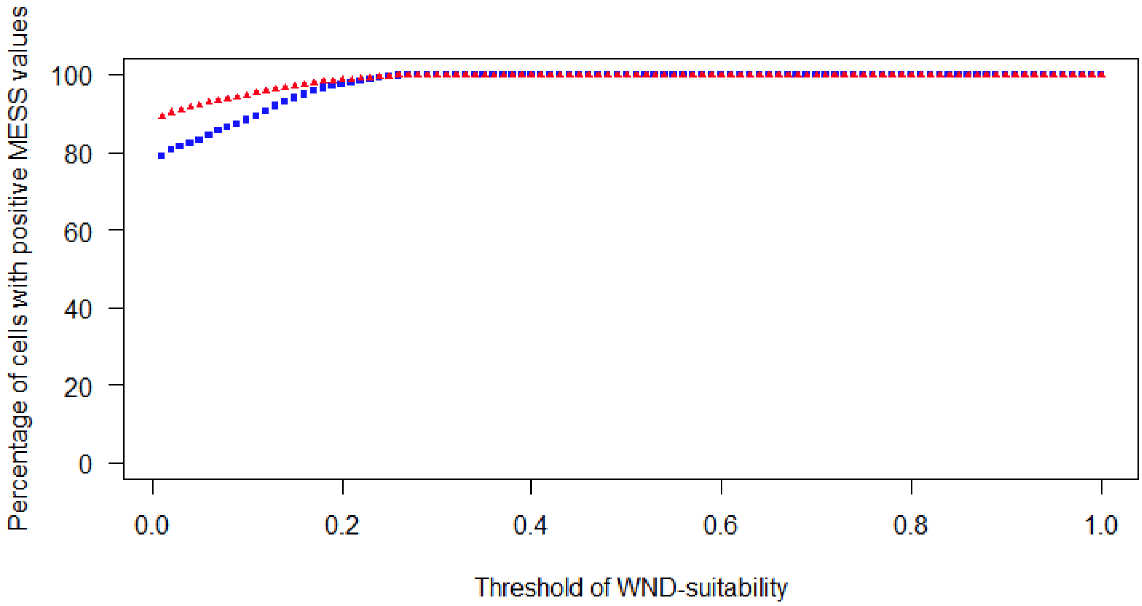


**Figure S9.** Results of the multivariate environmental similarity surface (MESS) analysis on the two environmental predictor variables herein used, MAST and precipitation. Positive values represent sites where no variable falls outside the range of environments observed in the reference set. Blue rectangles depict recent climatic conditions (1970-2000) while red triangles depict future conditions (2061-2080). The percentage of positive values reached 100% for WND-suitability > 0.27.

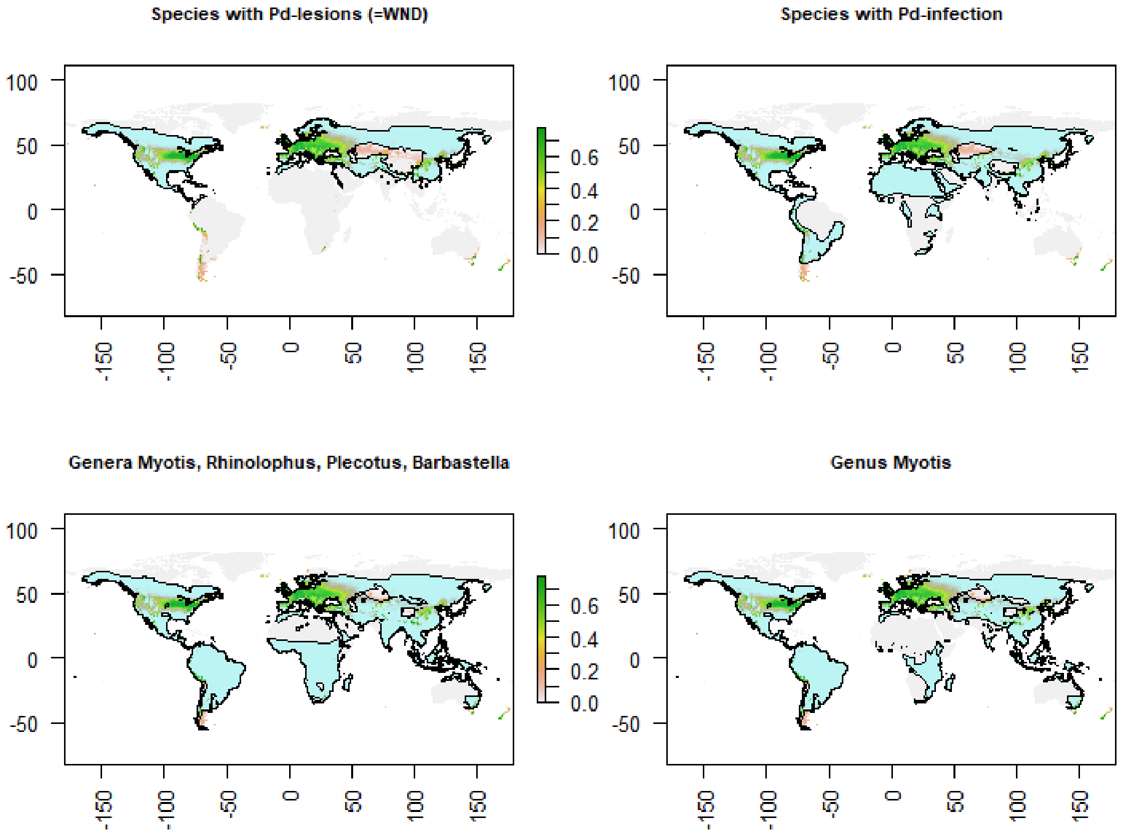


**Figure S10.** Maps of the world overlaying species-groups distribution for which we have some information about WND susceptibility and WND-suitability (predicted probability of WND occurrence) for recent climatic conditions (1970-2000). The species-groups mapped were: 1) top left panel, all species that have been recorded with WND (histopathological lesions confirmed), 2) top right panel, all species that have been recorded with *P. destructans* infection, 3) bottom left panel, all species in the bat genera included in the model (*Myotis, Rhinolophus, Plecotus, Barbastella*), 4) bottom right panel, all species in the bat genus classically harbouring the highest *P. destructans* load and lesions (i.e. *Myotis*). See main text for further details. Bat species distributions were recovered from the IUCN website as shape files (https://www.iucnredlist.org/; accessed 20.07.2023).

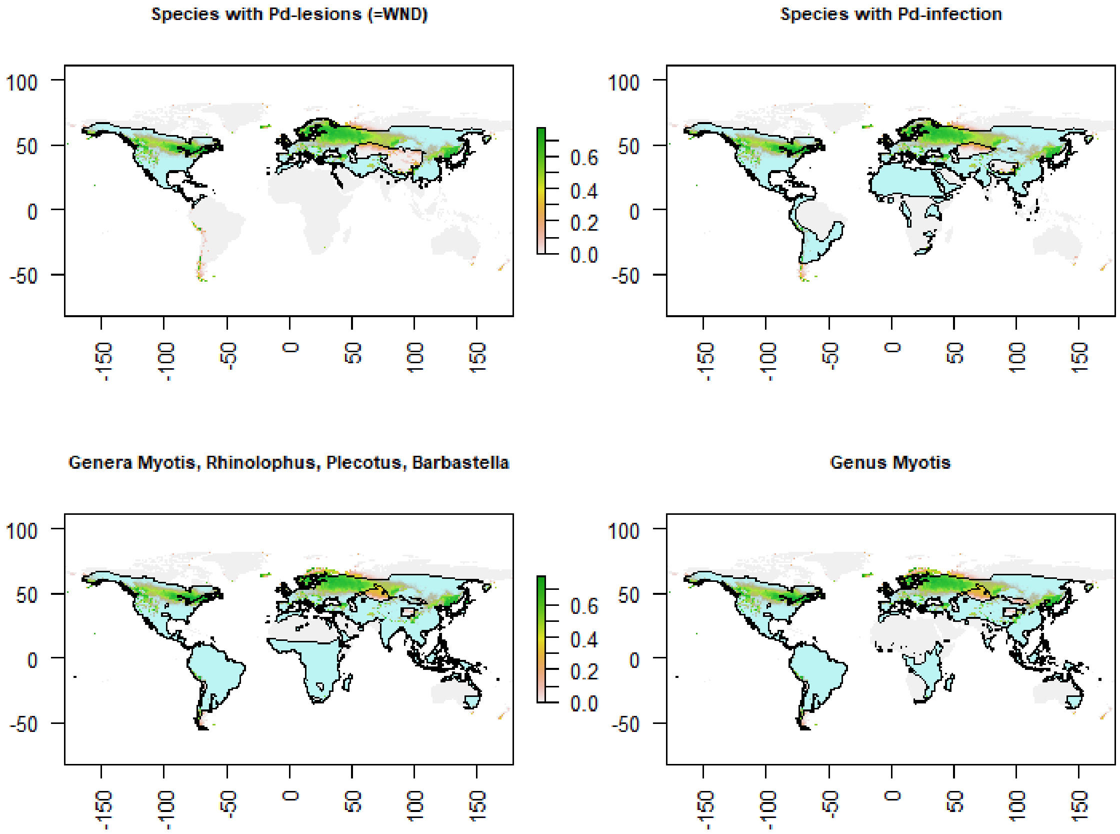


**Figure S11.** Maps of the world overlaying species-groups distribution for which we have some information about WND susceptibility and WND-suitability (predicted probability of WND occurrence) for future climatic conditions (2061-2080). Description of panels as per Fig. S10.
